# Nucleobase Methylation Enhances SARS-CoV-2 Chain Terminator Evasion of Exonuclease Proofreading

**DOI:** 10.1101/2024.09.20.614085

**Authors:** Lin Yang, Rory Cunnison, Carmen Ka Man Tse, Tiantian Xu, Buyu Zhang, Rick Xinzhou Xu, Benny Zhibin Liang, Xunyu Zhou, Danielle Groves, Adrian Deng, Jeremy R Keown, Daqi Yu, Mingyuan Li, Yuanliang Zhai, Billy Wai-Lung Ng, Lu Zhang, Nicole C Robb, Peter Pak-Hang Cheung

**Author notes:** These authors contributed equally to this work (Lin Yang, Rory Cunnison, Carmen Ka Man Tse, Tiantian Xu).

## Abstract

Inhibition of the SARS-CoV-2 RNA-dependent RNA polymerase (RdRp) by nucleoside analogues (NuAs) represents a key antiviral strategy. However, their efficacy is fundamentally limited by the viral proofreading exoribonuclease nsp14, which excises misincorporated NuAs from nascent viral RNA. To identify NuAs capable of achieving chain termination while resisting nsp14-nsp10-mediated excision, we sequentially assessed a panel of nucleotide analogues encompassing ribose 2′, 3′, and nucleobase modifications for RdRp chain termination and nsp14-nsp10 ExoN resistance. Among the compounds assessed, 5-methyl-3′-dUTP was identified as the most promising candidate, combining potent immediate chain termination with superior ExoN resistance. The 5-methyl modification confers substantial resistance to nsp14-nsp10-mediated proofreading excision, notably outperforming its unmodified counterpart 3′-dUTP. RdRp also exhibited improved incorporation efficiency for this analogue. RNA chains terminated with 5-methyl-3′-dUTP exhibit barely detectable primer extension even following nsp14-nsp10-mediated exonucleolytic cleavage. The superior chain termination and ExoN resistance of 5-methyl-3′-dUTP were independently corroborated using an smFRET assay. Mechanistically, molecular dynamics simulations suggest that this enhanced ExoN resistance arises from destabilization of F146 loop stacking in the nsp14-nsp10 active site. Collectively, these findings establish 5-methyl-3′-dUTP as a promising antiviral lead compound and provide a structural framework for designing next-generation nucleoside analogues capable of evading coronavirus proofreading surveillance.

## 1. Introduction

Coronavirus outbreaks have repeatedly highlighted the pandemic threat posed by this virus family, with SARS CoV 2, the causative agent of coronavirus disease 2019 (COVID 19), representing the most consequential outbreak in recent history. This virus has claimed millions of lives and continues to circulate globally through successive waves driven by immune-evading variants of concern (VOCs).^[1–3]^ Although effective vaccines have substantially reduced disease severity and mortality,^[4,5]^ their effectiveness is fundamentally constrained by antigenic drift, waning immunity, and the immunological vulnerability of high-risk populations, which vaccine-based approaches alone cannot fully address.^[6,7]^ More broadly, the recurring emergence of novel coronaviruses underscores that SARS CoV 2 is unlikely to be the last pathogen of its kind to threaten global health. Effective, broad spectrum antiviral therapeutics targeting conserved viral machinery therefore represent an essential complement to vaccine-based strategies, and a durable component of pandemic preparedness against current and future coronavirus threats.^[8]^

The RNA dependent RNA polymerase (RdRp) domain of SARS CoV 2 nsp12 represents a central target for antiviral drug development.^[9,10]^ The catalytically active holoenzyme comprises nsp12 and two accessory cofactors, nsp7 and nsp8, and orchestrates viral genome replication and transcription.^[11,12]^ The structural and functional conservation of the RdRp across coronaviruses, combined with its essential role in viral replication, positions it as an ideal target for developing broad spectrum antivirals.^[13]^ Nucleoside analogues (NuAs) constitute a well established class of RdRp inhibitors that exploit the viral polymerase by mimicking endogenous nucleoside substrates.^[14,15]^ Following cellular uptake, these prodrugs undergo sequential phosphorylation by host kinases to generate their active 5′ triphosphate forms (NuA TPs), which then compete with natural nucleoside triphosphates (NTPs) for incorporation into nascent viral RNA chains.^[14,15]^ Upon incorporation, NuAs can inhibit viral replication through diverse mechanisms, including chain termination and lethal mutagenesis that drives the viral population towards error catastrophe.^[16,17]^

Several nucleoside analogues have received emergency use authorization or demonstrated clinical promise for COVID 19 treatment, yet each is hampered by distinct limitations. Remdesivir, an adenosine analogue bearing a 1′ cyano modification, acts as a delayed chain terminator.^[16]^ While initial formulations required intravenous administration, orally bioavailable prodrugs have since been developed, including VV116 and obeldesivir.^[18,19]^ Remdesivir’s clinical efficacy remains modest in certain settings, likely reflecting multiple factors including the critical importance of early administration. Nevertheless, susceptibility to excision by the viral proofreading machinery constitutes a measurable biochemical constraint, as ExoN-deficient coronavirus mutants exhibit markedly enhanced remdesivir sensitivity.^[20–22]^ Molnupiravir, an orally bioavailable prodrug of N4 hydroxycytidine, induces viral error catastrophe through lethal mutagenesis, offering convenient administration but demonstrating only moderate efficacy and raising concerns regarding potential mutagenic effects on host cell DNA.^[23–25]^ Other candidates, including favipiravir and sofosbuvir, have shown limited effectiveness against SARS CoV 2.^[26–28]^ Among chain-terminating nucleoside analogues, susceptibility to excision by the viral nsp14-nsp10 proofreading complex represents a critical and unresolved vulnerability that has not been adequately addressed by any approved therapeutic to date.^[29,30]^ While alternative strategies such as mutagenesis-based inhibition have been explored, their own limitations underscore the broader challenge of developing nucleoside-based antivirals against coronaviruses with robust proofreading capacity.

Central to this challenge is a unique evolutionary adaptation that distinguishes coronaviruses from most other RNA viruses: a 3′ to 5′ exoribonuclease (ExoN) proofreading activity encoded by nsp14, which functions as a heterodimeric complex with its cofactor nsp10.^[29,31,32]^ This proofreading machinery recognizes and excises misincorporated nucleotides from the 3′ terminus of nascent viral RNA, including many nucleoside analogues incorporated by the RdRp, thereby attenuating their antiviral efficacy.^[22,33]^ The functional importance of this activity is underscored by genetic studies demonstrating that ExoN inactivation results in an approximately 15 fold increase in mutation frequency throughout the coronavirus genome, and that ExoN deficient mutants exhibit markedly increased susceptibility to numerous nucleoside analogues that are otherwise ineffective against wild type virus.^[34–36]^ Together, these findings establish the nsp14 nsp10 exonuclease complex as a critical barrier to nucleoside-based antiviral development.

Circumventing this barrier requires nucleoside analogues that satisfy two independent yet equally essential properties: efficient chain termination to block viral RNA synthesis, and robust resistance to excision by the nsp14 nsp10 complex to ensure durable inhibition.^[37]^ Structural modifications to the nucleotide scaffold offer a rational strategy to address both criteria. Modifications at the 3′ and 2′ positions of the ribose have been shown to confer chain termination activity through distinct mechanisms: 3′-deoxy substitutions act by eliminating the 3′-OH required for chain elongation, while 2′-modifications such as the 2′-methyl-2′-fluoro substitution in sofosbuvir sterically block incoming nucleotides, establishing ribose modification as a validated strategy for RdRp inhibition.^[15,38]^ However, despite the clear therapeutic rationale, a systematic framework for evaluating nucleotide scaffold modifications based on sequential assessment of chain termination activity followed by ExoN resistance remains lacking. The structural determinants governing ExoN resistance, in particular how specific nucleotide scaffold modifications disrupt nsp14-nsp10 activity, have not been fully elucidated, representing a fundamental gap that impedes the rational design of proofreading-resistant nucleoside analogues against coronaviruses.

In this study, we systematically evaluated a panel of structurally diverse nucleotide analogues encompassing modifications at the ribose 2′ position, 3′ position, and nucleobase, with the explicit goal of identifying compounds capable of satisfying both chain termination and proofreading resistance criteria. Among the candidates examined, 5 methyl 3′ dUTP stood out as the most promising inhibitor, functioning as a potent immediate chain terminator while demonstrating greater resistance to nsp14-nsp10-mediated excision than its unmodified counterpart 3′-dUTP. In addition, the viral RdRp showed superior incorporation efficiency for 5 methyl 3′ dUTP relative to 3′ dUTP. RNA chains terminated with 5 methyl 3′ dUTP remain highly resistant to further elongation even following nsp14-nsp10-mediated exonucleolytic cleavage and supplementation with the correct nucleotide. The superior chain termination and ExoN resistance of 5 methyl 3′ dUTP relative to 3′ dUTP were independently corroborated using smFRET assays, which monitor RNA conformational changes at the single-molecule level to report on both RdRp-mediated extension and nsp14-nsp10-mediated excision. Structural insights from MD simulations attributed this enhanced ExoN resistance to disruption of F146 loop stacking interactions within the nsp14-nsp10 active site. Collectively, this work establishes a sequential evaluation framework for screening nucleoside analogues against dual criteria of chain termination and ExoN resistance, and identifies 5 methyl 3′ dUTP as a promising lead compound for antiviral drug development, advancing our mechanistic understanding of how nucleotide scaffold modifications govern proofreading resistance. These results further offer a structural basis for the rational design of nucleoside analogues capable of circumventing the proofreading surveillance conserved across coronaviruses.

## 2. Results

### 2.1. Screening of Nucleoside Analogue Triphosphates (NuA-TPs) for Chain Termination Activity

To identify potential inhibitors of the SARS-CoV-2 RNA-dependent RNA polymerase (RdRp), we investigated a panel of nucleoside analogue triphosphates (NuA-TPs), including 2’-OMe-NTP, 2’-fluoro-dNTP, 3’-dNTP, 3’-(O-propargyl)-GTP, 5-methyl-3’-dUTP, and 6-aza-UTP, whose chemical structures are shown in **Figure 1**. We evaluated their ability to act as chain terminators during RNA synthesis by the viral RdRp complex using RNA extension assays. The LS2U (20nt) primer labeled with FAM at the 5’ end was used in these assays. For GTP and UTP analogue evaluation, the LS15 (40nt) template was paired with FAM-LS2U (**Figure 2A**), while the LS41-GU (41nt) template combined with FAM-LS2U was used for CTP and ATP analogue evaluation (**Figure 2D**).

**Figure 1.**
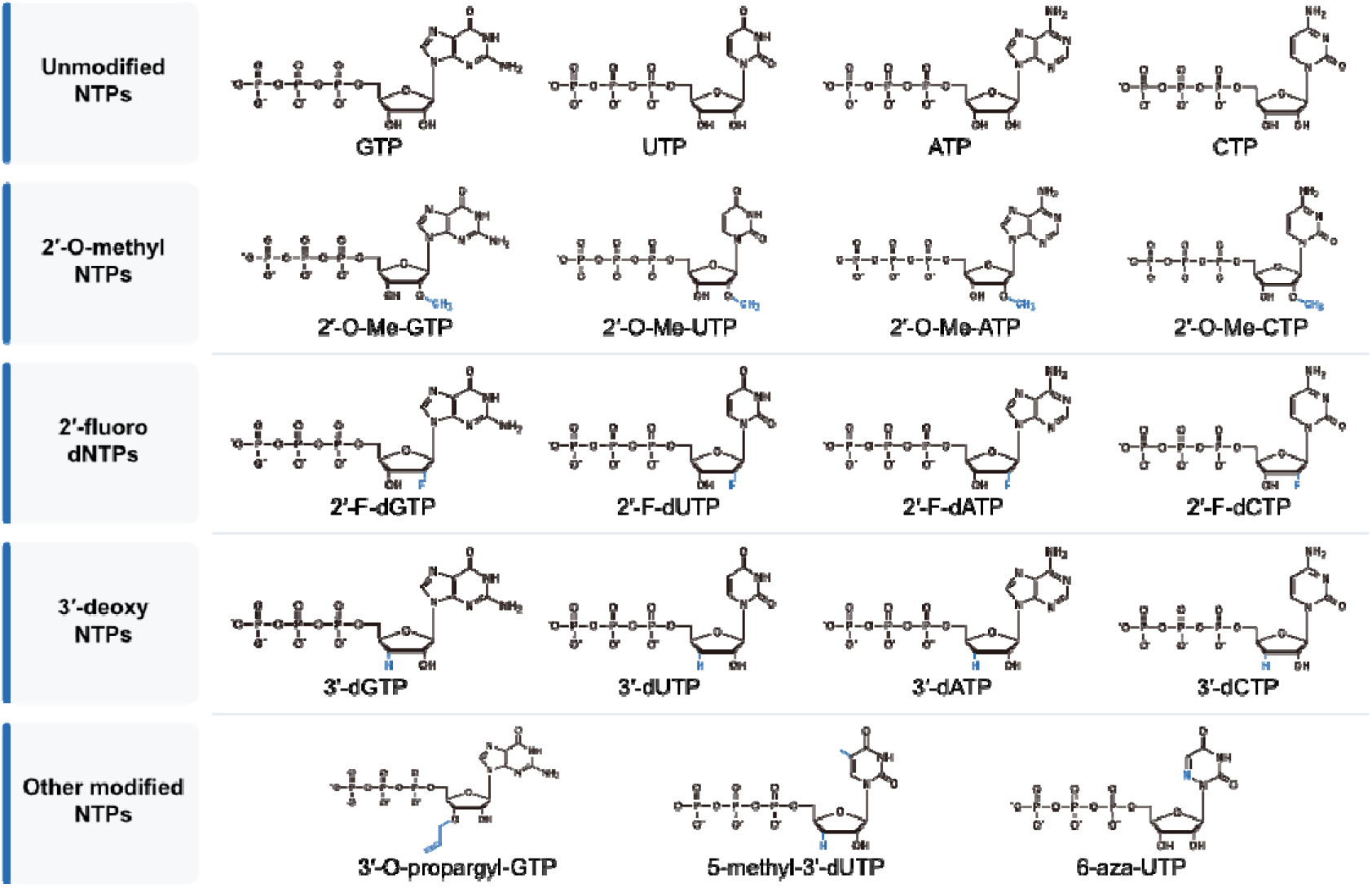
The chemical structures of the NuA-TPs employed in this study. All structures are drawn using a consistent phosphate protonation/charge convention, with counterions omitted, and retain the native D-ribose stereochemistry. NTP, nucleoside triphosphate; OMe, methoxy; F, fluoro; d, deoxy.

**Figure 2.**
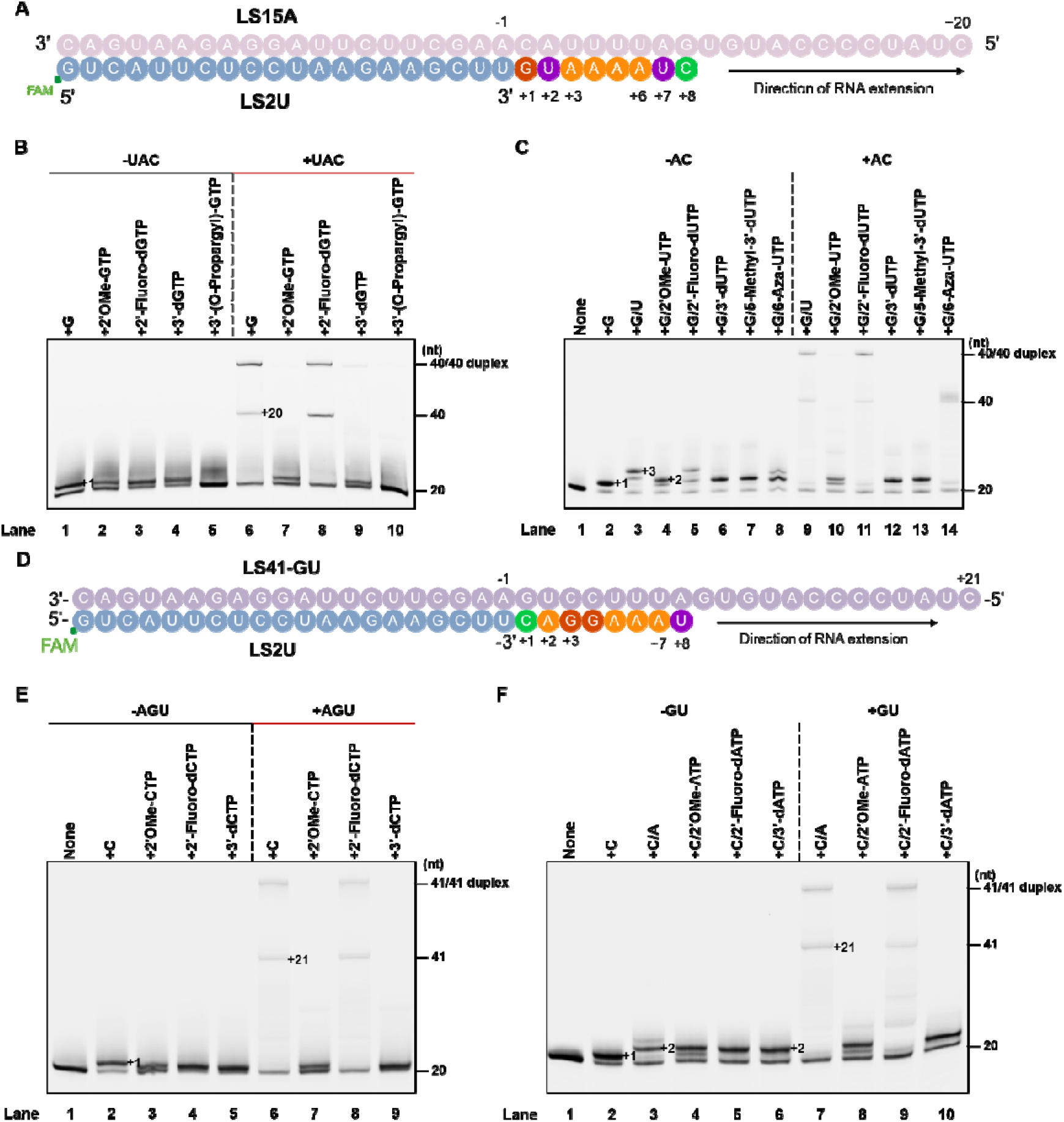
Evaluation of nucleoside analogue triphosphates (NuA-TPs) as chain terminators for SARS-CoV-2 RdRp. (A) RNA scaffolds used for GTP and UTP analogue assays. The experiment utilized the LS15A RNA template (40 nucleotides) paired with the LS2U RNA primer (20 nucleotides, 5’ fluorescently labeled).^[39]^ This scaffold was used for the RNA extension assays shown in panels (B) and (C) to test the chain termination effect of GTP and UTP analogues. (B) Gel electrophoresis results for GTP analogue chain termination. Assessment of the chain termination effect of various GTP analogues by SARS-CoV-2 RdRp on the LS15A/LS2U scaffold. (C) Gel electrophoresis results for UTP analogue chain termination. Assessment of the chain termination effect of various UTP analogues by SARS-CoV-2 RdRp on the LS15A/LS2U scaffold. (D) RNA scaffolds used for CTP and ATP analogue assays. The LS41-GU RNA template (41 nucleotides) paired with the LS2U RNA primer (20 nucleotides, 5’ fluorescently labeled) was used for the RNA extension assays shown in panels (E) and (F) to test the chain termination effect of CTP and ATP analogues. (E) Gel electrophoresis results for CTP analogue chain termination. Assessment of the chain termination effect of various CTP analogues by SARS-CoV-2 RdRp on the LS41-GU/LS2U scaffold. (F) Gel electrophoresis results for ATP analogue chain termination. Assessment of the chain termination effect of various ATP analogues by SARS-CoV-2 RdRp on the LS41-GU/LS2U scaffold.

As shown in **Figure 2B**, 2’-OMe-GTP, 2’-fluoro-dGTP, and 3’-dGTP were incorporated into the LS2U primer by RdRp, as indicated by the appearance of +1 bands comparable to those of the natural GTP control in the absence of UTP, ATP, and CTP (**Figure 2B**, lanes 1-4). In contrast, 3’-(O-propargyl)-GTP remained at the primer position, indicating that it was not incorporated by RdRp (**Figure 2B**, lane 5). Upon addition of UTP, ATP, and CTP, 2’-fluoro-dGTP-incorporated RNA proceeded to full-length products (**Figure 2B**, lane 8), similar to the natural GTP control (**Figure 2B**, lane 6), indicating that 2’-fluoro-dGTP does not terminate SARS-CoV-2 RdRp-mediated synthesis. Conversely, 2’-OMe-GTP and 3’-dGTP predominantly remained as +1 bands even in the presence of all three NTPs (**Figure 2B**, lanes 7 and 9), demonstrating that both analogues function as immediate chain terminators.

We further assessed the chain termination activity of UTP analogues bearing distinct structural modifications. The RNA LS15A template was designed such that GTP incorporation precedes UTP incorporation. In the absence of ATP and CTP, we observed two distinct patterns (**Figure 2C**). Similar to the natural UTP control, 2’-fluoro-dUTP and 6-aza-UTP generated a +3 band, indicating that after their incorporation at the +2 position, the RdRp misincorporated an additional nucleotide (**Figure 2C**, lanes 3, 5, and 8). In contrast, 2’-OMe-UTP, 3’-dUTP, and 5-methyl-3’-dUTP produced the +2 band, showing they halted synthesis immediately after incorporation (**Figure 2C**, lanes 4, 6, and 7). Upon addition of ATP and CTP, the RNA incorporated with 2’-fluoro-dUTP and 6-aza-UTP was extended to full-length products (**Figure 2C**, lanes 11 and 14), similar to the UTP control (**Figure 2C**, lane 9), confirming they are non-terminating analogues. In contrast, reactions with 2’-OMe-UTP, 3’-dUTP, and 5-methyl-3’-dUTP remained as +2 position in the presence of ATP and CTP (**Figure 2C**, lanes 10, 12, and 13), confirming their function as immediate chain terminators.

Similar results were obtained for CTP analogues. As demonstrated in **Figure 2E**, 2’-OMe-CTP, 2’-fluoro-dCTP, and 3’-dCTP were incorporated into the LS2U primer by RdRp, producing +1 bands in the absence of ATP, GTP, and UTP (**Figure 2E**, lanes 3-5). Upon addition of the missing NTPs, 2’-fluoro-dCTP-incorporated RNA proceeded to full-length products (**Figure 2E**, lane 8), similar to the natural CTP control (**Figure 2E**, lane 6), indicating non-terminating behavior. However, 2’-OMe-CTP and 3’-dCTP predominantly remained as +1 bands in the presence of all three NTPs (**Figure 2E**, lanes 7 and 9), confirming their immediate terminating activity.

A consistent pattern was observed for ATP analogues. **Figure 2F** shows that 2’-OMe-ATP, 2’-fluoro-dATP, and 3’-dATP were incorporated into the LS2U primer by RdRp, producing +2 bands in the absence of GTP and UTP (**Figure 2F**, lanes 4-6). Upon addition of GTP and UTP, 2’-fluoro-dATP-incorporated RNA proceeded to full-length products (**Figure 2F**, lane 9), similar to the natural ATP control (**Figure 2F**, lane 7), indicating non-terminating behavior. Conversely, 2’-OMe-ATP and 3’-dATP remained as +2 bands in the presence of GTP and UTP (**Figure 2F**, lanes 8 and 10), demonstrating immediate chain termination.

Taken together, these results revealed distinct incorporation and termination profiles among the tested nucleoside analogues. 3’-(O-propargyl)-GTP could not be incorporated by RdRp, while all other analogues were successfully incorporated. Among the incorporated analogues, 2’-fluoro-dNTP and 6-aza-UTP allowed continued RNA synthesis to full-length products, indicating they do not function as chain terminators. In contrast, 2’-OMe-NTP, 3’-dNTP, and 5-methyl-3’-dUTP functioned as immediate chain terminators, effectively blocking SARS-CoV-2 RNA synthesis upon incorporation.

### 2.2. Resistance of Terminated RNA to Viral Proofreading Exonuclease (nsp14-nsp10)

Coronaviruses possess a proofreading exonuclease (ExoN) comprising the nsp14-nsp10 complex, which can remove misincorporated nucleotides and attenuate the efficacy of antiviral nucleoside analogues.^[30]^ Having identified 2’-OMe-NTP, 3’-dNTP, and 5-methyl-3’-dUTP as immediate chain terminators, we next evaluated their resistance to nsp14-nsp10-mediated proofreading excision. The experimental workflow is illustrated in **Figure 3A**. We used the FAM-LS2U primer paired with the LS15A template for these assays (**Figure 2A**). The protocol involved a sequential three-step process: first, the RNA template/primer duplex was incubated with SARS-CoV-2 RdRp, the corresponding NTP analogue, and the three natural NTPs at 37°C for 10 minutes to enzymatically generate analogue-terminated RNA products. Subsequently, nsp14-nsp10 was added to the reaction and incubated for an additional 10 minutes at 37°C to assess exonucleolytic cleavage. Finally, the products were analyzed by denaturing PAGE.

**Figure 3.**
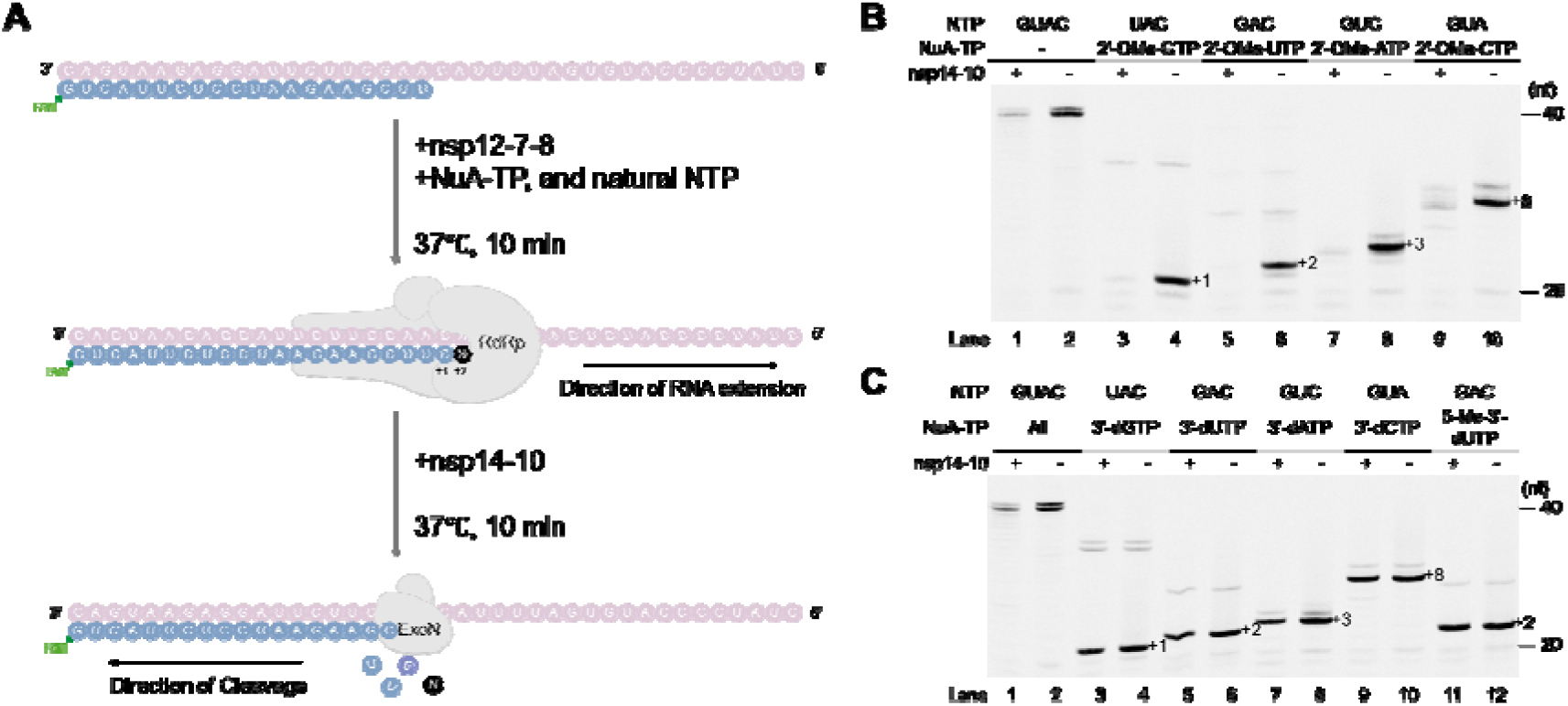
RdRp extension-ExoN cleavage assays of 2’-OMe-NTP, 3’-dNTP, and 5-Me-3’-dUTP. (A) Experimental scheme for extension-cleavage assays. Schematic diagram illustrating the multi-step protocol for generating analogue-incorporated RNA substrates by RdRp extension, followed by incubation with the SARS-CoV-2 nsp14-nsp10 complex to test for exonuclease (ExoN) removal. Results are shown in panels (B) and (C). (B) Cleavage of 2’-OMe-ribonucleoside-incorporated RNA substrates. Gel electrophoresis results showing the removal of RNA substrates containing 2’-OMe-ribonucleoside analogues by the SARS-CoV-2 nsp14-nsp10 proofreading complex. (C) Cleavage of 3’-deoxyribonucleoside and 5-Me-3’-deoxyuridine-incorporated RNA substrates. Gel electrophoresis results showing the resistance to removal for RNA substrates containing 3’-deoxyribonucleosides or 5-Me-3’-deoxyuridine by the SARS-CoV-2 nsp14-nsp10 proofreading complex.

The results revealed striking differences in exonuclease susceptibility among the tested analogues. As shown in **Figure 3B**, RNA chains terminated with 2’-OMe-NTP demonstrated high susceptibility to cleavage by the nsp14-nsp10 complex relative to the natural NTP control. The termination effects of 2’-OMe-NTP in the absence of nsp14-nsp10 are shown in lanes 4, 6, 8, and 10, while the corresponding cleavage patterns following nsp14-nsp10 treatment are displayed in lanes 3, 5, 7, and 9 (**Figure 3B**). The majority of RNA terminated with 2’-OMe-NMP was efficiently cleaved by the proofreading exonuclease, indicating that this modification alone is insufficient to evade the viral defense mechanism. In contrast, RNA chains terminated with either 3’-dNTP or 5-methyl-3’-dUTP exhibited remarkable resistance to exonucleolytic removal compared to the natural NTP control. As demonstrated in **Figure 3C** (lanes 3, 5, 7, 9), these analogue-terminated RNAs showed only minimal cleavage by the nsp14-nsp10 complex compared to the control conditions (lanes 4, 6, 8, 10). Such pronounced resistance to proofreading activity suggests that both 3’-dNTP and 5-methyl-3’-dUTP possess the critical ability to evade this key viral defense mechanism, suggesting their potential as candidates for further antiviral development.

### 2.3. 5-Methyl-3’-dUMP Confers Superior Resistance to Exonuclease Cleavage

Given that both 3’-dNTP and 5-methyl-3’-dUTP demonstrated resistance to nsp14-nsp10-mediated excision (**Figure 3B** and **3C**), we next compared the cleavage susceptibility of RNA substrates terminated with 3′-dUMP versus 5-methyl-3′-dUMP to determine whether the 5-methyl modification on the nucleobase further enhances this resistance, using both concentration-dependent and time-course analyses. Using the same FAM-LS2U/LS15A template-primer system, the RNA template/primer duplex was first incubated with SARS-CoV-2 RdRp, GTP, and UTP or the corresponding UTP analogue (3’-dUTP, or 5-methyl-3’-dUTP) at 37°C for 10 minutes to enzymatically generate UMP-, 3’-dUMP-, or 5-methyl-3’-dUMP-terminated RNA products.

For the concentration-dependent analysis, serial dilutions of nsp14-nsp10 were added to the respective reactions, and the products were analyzed by denaturing PAGE. The results demonstrated that 5-methyl-3’-dUMP-terminated RNA exhibited consistently greater resistance to nsp14-nsp10 cleavage compared to 3’-dUMP-terminated RNA in a concentration-dependent manner (**Figure 4A** and **4C**). Quantification confirmed that at the highest nsp14-nsp10 concentration tested (300 nM), 3′-dUMP-terminated RNA showed a relative band intensity of nearly 20%, whereas 5-methyl-3′-dUMP-terminated RNA retained approximately 40% (Figure 4C), with significant differences observed at all concentrations tested. In the time-course experiment, a fixed concentration of nsp14-nsp10 was added to each reaction and incubated for 0, 5, 10, 20, and 40 minutes at 25°C before analysis by denaturing PAGE. Consistent with the concentration-dependent results, 5-methyl-3’-dUMP-terminated RNA showed markedly enhanced resistance to nsp14-nsp10 cleavage over time compared to 3′-dUMP-terminated RNA (**Figure 4B** and **4D**). By the 40-minute time point, 3′-dUMP-terminated RNA was nearly completely degraded, whereas 5-methyl-3′-dUMP-terminated RNA maintained approximately 20% relative band intensity (Figure 4D), demonstrating sustained resistance to nsp14-nsp10-mediated excision throughout the time course.

**Figure 4.**
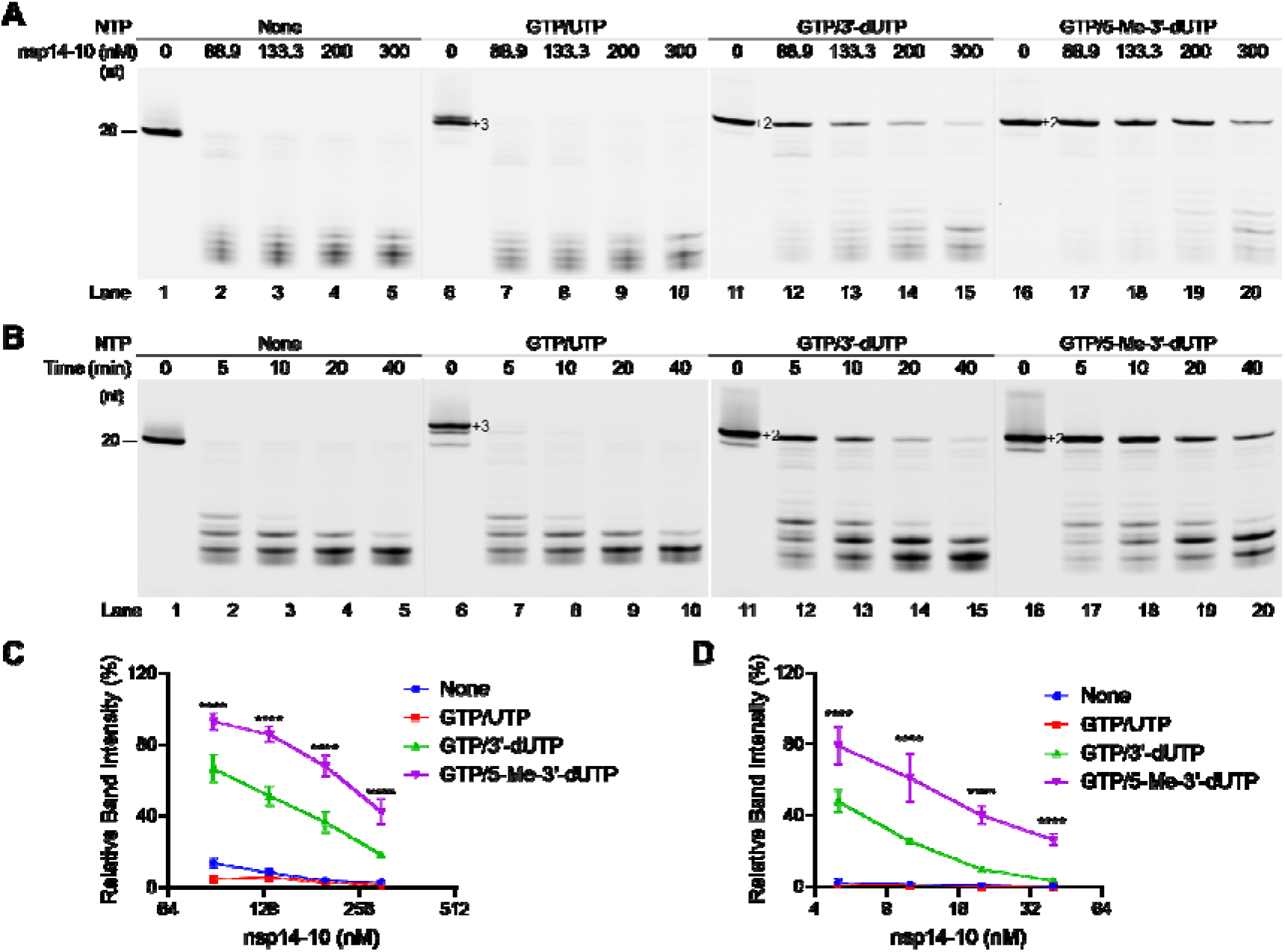
Differential exonucleolytic cleavage of 3’-dUMP and 5-Me-3’-dUMP RNA substrates by nsp14-nsp10. (A) Concentration-dependent cleavage assay of 3’-dUMP and 5-Me-3’-dUMP substrates. Representative gel image showing the cleavage of RNA substrates incorporating 3’-deoxyuridine (3’-dUMP residual) or 5-methyl-3’-deoxyuridine (5-Me-3’-dUMP residual) when incubated with increasing concentrations of the SARS-CoV-2 nsp14-nsp10 complex (ranging from 0 to 300 nM, 1.5-fold serial dilution). (B) Time-course cleavage assay of 3’-dUMP and 5-Me-3’-dUMP substrates. Representative gel image showing the time-dependent cleavage of RNA substrates incorporating 3’-dUMP or 5-Me-3’-dUMP residuals using a single, fixed concentration of the SARS-CoV-2 nsp14-nsp10 complex (150 nM). (C) Quantification of concentration-dependent cleavage. Quantification of the data presented in panel (A), showing the percentage of remaining uncleaved RNA substrate versus nsp14-nsp10 concentration. (D) Quantification of time-course cleavage. Quantification of the data presented in panel (B), showing the percentage of remaining uncleaved RNA substrate over time. Data are presented as means ± SD from three independent experiments (n = 3). *, **, ***, **** symbols denote the statistical significance of the difference between the 3’-dUMP group and the 5-Me-3’-dUMP group. Statistical significance is defined as: *p < 0.05, **p < 0.01, ***p < 0.001, ****p < 0.0001; N.S., not significant. Two-way ANOVA was used for two-factor multi-group comparisons.

The nsp14-nsp10 exonuclease cleavage assays (**Figure 4**) demonstrated that RNA substrates containing 5-methyl-3’-dUMP at the 3’ terminus are substantially more resistant to cleavage than those ending with 3’-dUMP. To independently corroborate this finding, we employed real-time fluorescence polarization (FP) analysis (**Figure S3**A and **S3**B), which monitors the rotational mobility of fluorescently labeled molecules to detect changes in molecular size. Upon addition of nsp14-nsp10, the FP signal of RNA substrates terminated with 5-methyl-3’-dUMP decreased more slowly compared to those ending with 3’-dUMP, indicating reduced exonuclease activity. These results confirmed that 5-methyl-3’-dUMP-terminated RNA serves as a poorer substrate for the nsp14-nsp10 exonuclease, consistent with a critical role of the 5-methyl modification in conferring enhanced resistance to nsp14-nsp10-mediated proofreading.

### 2.4. Incorporation Efficiency of 5-Methyl-3’-dUTP by RdRp

Having established the superior ExoN resistance of 5-methyl-3’-dUTP over 3’-dUTP, we next assessed whether the 5-methyl modification also affects incorporation efficiency by the RdRp, as this determines the analogue’s ability to compete with natural UTP for incorporation. We compared the incorporation efficiency of 5-methyl-3’-dUTP against 3’-dUTP by the SARS-CoV-2 RdRp using single nucleotide incorporation assays. In this experiment, primer-template complexes were incubated with increasing concentrations of each nucleotide analogue, and the extension of primers by a single nucleotide was monitored by denaturing PAGE. The results (**Figure 5**) revealed distinct incorporation profiles for each substrate. Natural UTP exhibited the highest incorporation efficiency with an EC50 (half-maximal effective concentration) of 47.88 nM, consistent with its role as the native substrate. Notably, 5-methyl-3’-dUTP demonstrated substantially improved incorporation compared to 3’-dUTP, with EC50 values of 106.7 nM and 750.8 nM, respectively, representing approximately a 7-fold enhancement in incorporation efficiency conferred by the 5-methyl modification. These findings suggest that the 5-methyl modification enhances recognition and utilization of this analogue by the viral polymerase.

**Figure 5.**
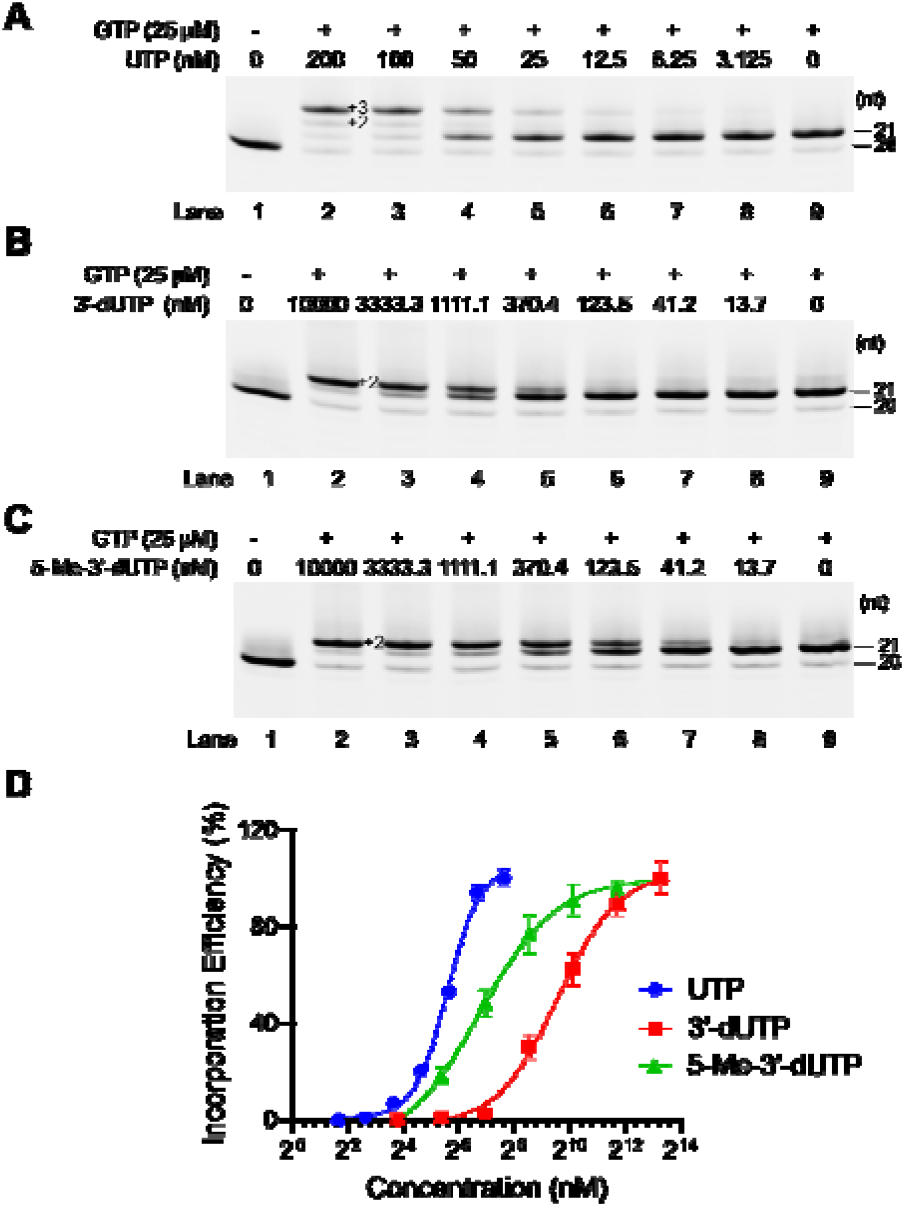
Determination of incorporation efficiency of 3’-dUTP and 5-methyl-3’-dUTP by SARS-CoV-2 RdRp. (A) Representative gel image for UTP incorporation. RNA extension reactions performed in the presence of increasing concentrations of the natural substrate UTP (ranging from 0 to 200 nM, 2-fold serial dilution). (B) Representative gel image for 3’-dUTP incorporation. RNA extension reactions performed in the presence of increasing concentrations of the analogue 3’-dUTP (ranging from 0 to 10 μM, 3-fold serial dilution). (C) Representative gel image for 5-methyl-3’-dUTP incorporation. RNA extension reactions performed in the presence of increasing concentrations of the analogue 5-methyl-3’-dUTP (ranging from 0 to 10 μM, 3-fold serial dilution). (D) Dose-response curves for UTP, 3’-dUTP, and 5-methyl-3’-dUTP. Dose-response curves were generated by quantifying the full extension products from panels (A), (B), and (C) across three independent experiments (n=3). The data are presented as means ± SD and were fitted using a four-parameter logistic model to determine incorporation efficiency.

### 2.5. RNA Chains Terminated with 3’-dUTP or 5-Methyl-3’-dUTP Cannot Be Rescued After Exonucleolytic Cleavage

Having established that 3’-dNTP and 5-methyl-3’-dUTP resist nsp14-nsp10-mediated excision, we next examined whether any residual cleavage could lead to rescue of viral RNA synthesis. The experimental workflow is illustrated in **Figure 6A**. The FAM-LS2U primer paired with the LS15A template was used for these cleavage-reextension assays. The protocol involved a two-step process: first, the RNA template/primer duplex was incubated with SARS-CoV-2 RdRp, ATP, CTP, GTP, and the corresponding NTP analogue (sofosbuvir-TP, 3’-dUTP, or 5-methyl-3’-dUTP) at 37°C for 20 minutes to generate analogue-terminated RNA products. Subsequently, nsp14-nsp10 and natural UTP were added to the reaction to assess whether residual exonucleolytic cleavage could rescue RNA synthesis.

**Figure 6.**
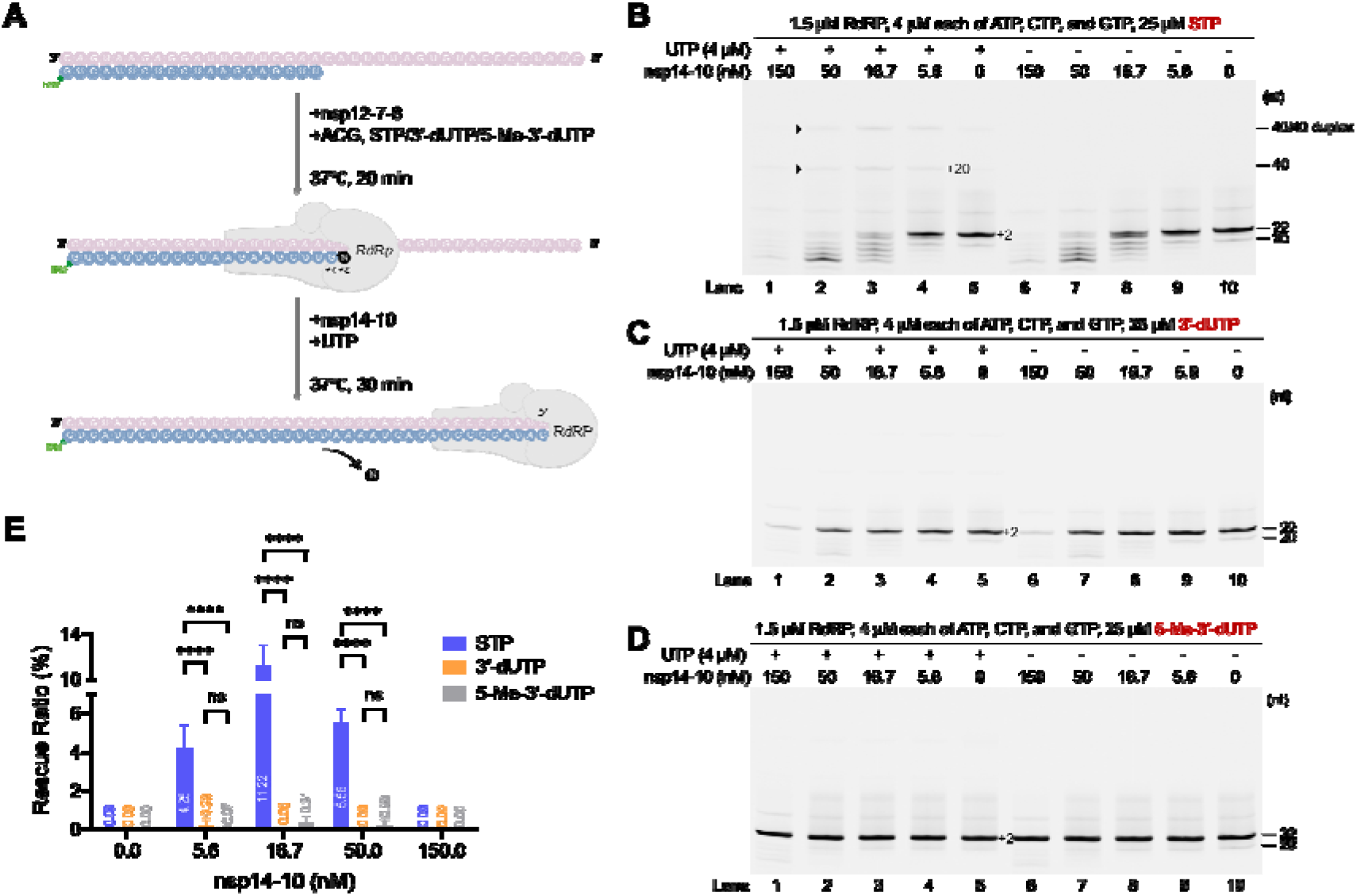
Cleavage-reextension assay to evaluate RNA substrate rescue. (A) Experimental workflow of the cleavage-reextension assay. Schematic diagram illustrating the sequential steps: (1) Nucleoside analogue (NuA-TP) incorporation by RdRp to create terminated RNA, (2) nsp14-nsp10 cleavage of the terminated product, and (3) RdRp-mediated reextension in the presence of natural nucleoside triphosphates (NTPs) to assess chain rescue potential. (B) Cleavage-reextension assay results for sofosbuvir-TP-terminated RNA. Representative gel image showing the rescue potential of RNA substrates terminated by the sofosbuvir analogue. (C) Cleavage-reextension assay results for 3’-dUTP-terminated RNA. Representative gel image showing the rescue potential of RNA substrates terminated by 3’-dUTP. (D) Cleavage-reextension assay results for 5-methyl-3’-dUTP-terminated RNA. Representative gel image showing the rescue potential of RNA substrates terminated by 5-methyl-3’-dUTP. (E) Quantification of RNA substrate rescue efficiency. Quantification of the full-length reextension products from panels (B), (C), and (D). Data are presented as means ± SD from three independent experiments (n = 3). *, **, ***, **** symbols denote the statistical significance of the difference between the 3’-dUMP group and the 5-Me-3’-dUMP group. Statistical significance is defined as: *p < 0.05, **p < 0.01, ***p < 0.001, ****p < 0.0001; N.S., not significant. Two-way ANOVA was used for two-factor multi-group comparisons.

Sofosbuvir-TP, a known chain terminator whose terminated RNA can be rescued by nsp14-nsp10-mediated cleavage followed by re-extension,^[39]^ was included as a positive control for chain rescue. As shown in **Figure 6B**, the sofosbuvir-TP-terminated RNA was efficiently cleaved by nsp14-nsp10 and could be subsequently extended upon addition of natural UTP, demonstrating successful rescue of the terminated RNA chain. In stark contrast, RNA chains terminated with either 3’-dUTP or 5-methyl-3’-dUTP showed markedly different behavior. Despite treatment with nsp14-nsp10 and supplementation with natural UTP, these analogue-terminated RNA chains remained refractory to extension (**Figure 6C**-**6D**). Quantification of rescue ratios confirmed this observation: the rescue ratio of sofosbuvir-TP-terminated RNA reached approximately 11% at 16.7 nM nsp14-nsp10, whereas 3′-dUTP- and 5-methyl-3′-dUTP-terminated RNA maintained rescue ratios near 0% across all concentrations tested (Figure 6E). These results indicate that 3’-dUTP and 5-methyl-3’-dUTP effectively resist nsp14-nsp10-mediated excision and prevent subsequent rescue of RNA synthesis by RdRp, suggesting their potential as promising scaffolds for nucleoside analogue design.

### 2.6. Single-Molecule FRET Assay Confirms Superior Performance of 5-Methyl-3’-dUTP

To independently corroborate our biochemical findings using an orthogonal approach, we employed a solution-based single-molecule Förster resonance energy transfer (smFRET) assay, previously developed to monitor conformational changes in the RNA during SARS-CoV-2 RdRp-mediated RNA extension and inhibition.^[40]^ We used a 43-nt RNA template engineered to form a Cy3-labeled hairpin, annealed to a shorter primer carrying a Cy5 label (**Figure S4**A). This duplex positions the dyes close together, producing a high FRET efficiency (E) signal (E∼0.85) (**Figure S4**B top panel, **S4**C). Upon RdRp binding and nucleotide incorporation, the primer extends and disrupts the hairpin, separating the dyes and resulting in a low FRET signal (E∼0.15) (**Figure S4**B lower panel, **S4**C).

Our initial assay design relied on multiple ATP incorporation to ensure initial opening of the hairpin. To test the specific addition of UTP, 3’-dUTP or 5-Methyl-3’-dUTP, we therefore modified the template sequence so that UTP or its analogues were incorporated first **(Figure S4D**). Similarly to the original template, RNA only provided a high FRET signal (E∼0.8) (**Figure S4**E top panel, **S4**F), although we note that this FRET distribution is wider than that of the original design, possibly due to less efficient closing of the hairpin or the hairpin being bound in several different registers. Nevertheless, upon addition of RdRp and 300 µM NTPs, we observed a distinct low FRET population (E∼0.2), indicative of efficient RNA extension (**Figure S4**E middle panel, **S4**F). Full conversion of the hairpin to open, extended RNA (E∼0.2) was observed even in the presence of a lower (10 µM) concentration of NTPs (**Figure S4**E bottom panel, **S4**F).

Next, we applied this approach to assess incorporation of 5-Methyl-3’-dUTP or 3’-dUTP, in the presence of either a high (300 µM) or low (10 µM) concentration of competing natural UTP. We incubated RNA, RdRp, and increasing concentrations of 5-Methyl-3’-dUTP, alongside 300 µM of natural UTP and ATP, and observed that the RNA primer was almost fully extended at 50 µM 5-Methyl-3’-dUTP, whilst there was partial inhibition of extension with 150 µM 5-Methyl-3’-dUTP (**Figure 7A**). When the concentration of 5-Methyl-3’-dUTP was increased to 500 µM a large decrease in the low FRET population representing fully extended RNA was observed (E∼0.2); interestingly coinciding with a shift in the higher FRET population to an intermediate FRET value (E∼0.6), likely representing a partially open hairpin that could be partly extended RNA (**Figure 7A**). Significantly increased inhibition was observed with as little as 50 µM 5-Methyl-3’-dUTP when the concentration of competing natural UTP was reduced to 10 µM (**Figure 7B**). In contrast, when the activity of the RdRp in the presence of increasing concentrations of 3’-dUTP and high (300 µM) concentrations of natural UTP was assessed, we observed almost full extension of the RNA (i.e. no inhibition), even at 500 µM 3’-dUTP (**Figure 7C**). Partial inhibition of RdRp activity was observed with 50 and 150 µM 3’-dUTP when the concentration of competing natural UTP was reduced to 10 µM, with increased levels of inhibition observed with 500 µM 3’-dUTP (**Figure 7D**). Together, these results confirm the observations of the RdRp Enzymatic Activity Assay, further proving that while both 5-Methyl-3’-dUTP and 3’-dUTP are capable of preventing RdRp-mediated extension, 5-Methyl-3’-dUTP is more efficient at doing so.

**Figure 7.**
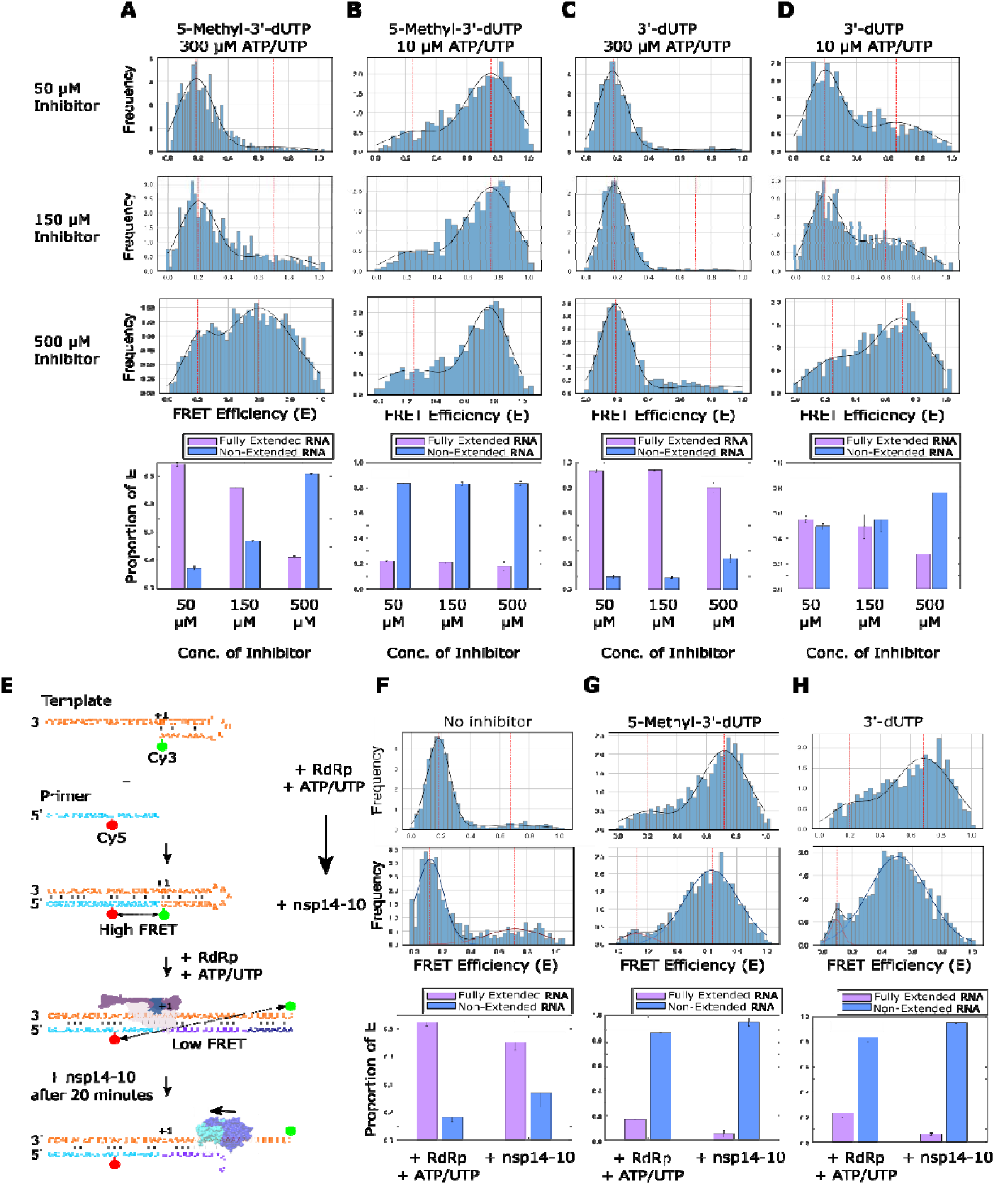
Inhibition of RNA extension with 5-Methyl-3’-dUTP and 3’-dUTP. (A-D) FRET efficiencies, E, for RNA conformations during extension by the RdRp in high (300 µM) or low (10 µM) concentrations of natural ATP/UTP, and increasing concentrations of 5-Methyl-3’-dUTP or 3’-dUTP. RNA was incubated with RdRp, either 300 or 10 µM ATP/UTP, and either 50, 150 or 500 µM inhibitor before being diluted for single-molecule FRET measurements. Quantification of the high and low FRET populations in panels A-D are shown as a proportion of the total FRET distributions by finding the probability density function. Error bars represent standard error of the Gaussian fit. (E) Schematic of experimental pipeline used to test the effect of nsp14-nsp10 addition to the single-molecule FRET assay. (F-H) FRET efficiencies, E, for RNA conformations during extension by the RdRp in the presence of 10 µM natural ATP/UTP and either no inhibitor, or 500 µM 5-Methyl-3’-dUTP or 3’-dUTP (top row). Nsp14-nsp10 was then added to the reactions after 20 minutes, after which further measurements were taken (middle row). Quantification of the high and low FRET populations in panels F-H are shown as a proportion of the total FRET distributions by finding the probability density function. Error bars represent standard error of the Gaussian fit.

Finally, we analysed the extension products of the RdRp in low natural UTP (10 µM) conditions after nsp14-nsp10 addition (**Figure 7E**), with either no nucleoside analogue present, or with 5-Methyl-3’-dUTP or 3’-dUTP present. As expected, we observed almost full extension of the primer in the absence of any inhibitor, which was almost fully retained when nsp14-nsp10 was added to the reaction after 20 minutes (**Figure 7F**). In the presence of 500 µM 5-Methyl-3’-dUTP or 3’-dUTP extension activity was fully or partially inhibited respectively, confirming our results above (**Figure 7G&H**; top row). When nsp14-nsp10 was added to the reaction after 20 minutes the proportion of non-extended RNA remained largely the same in both conditions, although we did observe a shift of the non-extended RNA population to intermediate FRET values for both inhibitors, suggesting an increase in a partially open RNA hairpin structure (**Figure 7G** and **7H**; bottom row). Together, these observations support the finding that both 5-Methyl-3’-dUTP and 3’-dUTP are resistant to cleavage by the nsp14-nsp10 exonuclease after being incorporated into the 3’ end of an RNA chain.

### 2.7. Molecular dynamics simulation suggested nucleobase 5-methylation enhances evasion of exonuclease proofreading by additional destabilization against F146 stacking

To obtain structural insight of the enhanced inhibition from the 5-methyl group modification, we performed molecular dynamics (MD) simulations on nsp14-nsp10-RNA complex containing 3’-dU and 5-Methyl-3’-dU, with U and 2’-O-Methyl-U included as controls. The simulation models were built from published cryo-EM SARS-CoV-2 nsp14-nsp10-RNA structure,^[41]^ with 10 replicas of 100 ns MD simulations generated for each system for analysis (see Supplementary Information).

We first characterized the catalytic active configurations in nsp14-nsp10-RNA complex. Among the catalytic residues, H268 of nsp14 serves as the critical general base to deprotonate the catalytic water during the phosphoryl-transfer reaction,^[41]^ which is conserved across all coronaviruses and critical for nsp14 ExoN activity and viral replication.^[42,43]^ Structurally, H268 encloses the active site in the presence of RNA substrate^[41]^ and is positioned to activate the Mg^2+^-bound catalytic water (Water_CAT_) in the solvated model (**Figure 8A**). Therefore, we first compared the total population of catalytic active configurations with intact contact between H268–Water_CAT_ among the four systems (**Figure 8B**). Our results showed that while the loss of 3’-OH reduced the number of active configurations from 54% (U) to 30% (3’-dU), the 5-methylation further decreased the number to 19% (5-Methyl-3’-dU), while 2’-O-Methyl-U showed a comparable population as 52% (2’-O-Methyl-U). The drastic effect of 3’-deoxy is consistent with early observation that 3’-OH is responsible for substrate recognition by the catalytic residue E92.^[41]^ In our MD simulations, the distance between the C3’ atom of terminal nucleotide and E92 showed a general increase for both 3’-dU and 5-Methyl-3’-dU when compared with U and 2’-O-Methyl-U (**Figure S5**). As a result, the loss of this interaction consistently destabilized the enclosed active site and led to a less stable Water_CAT_ binding and H268 loop closure.

**Figure 8.**
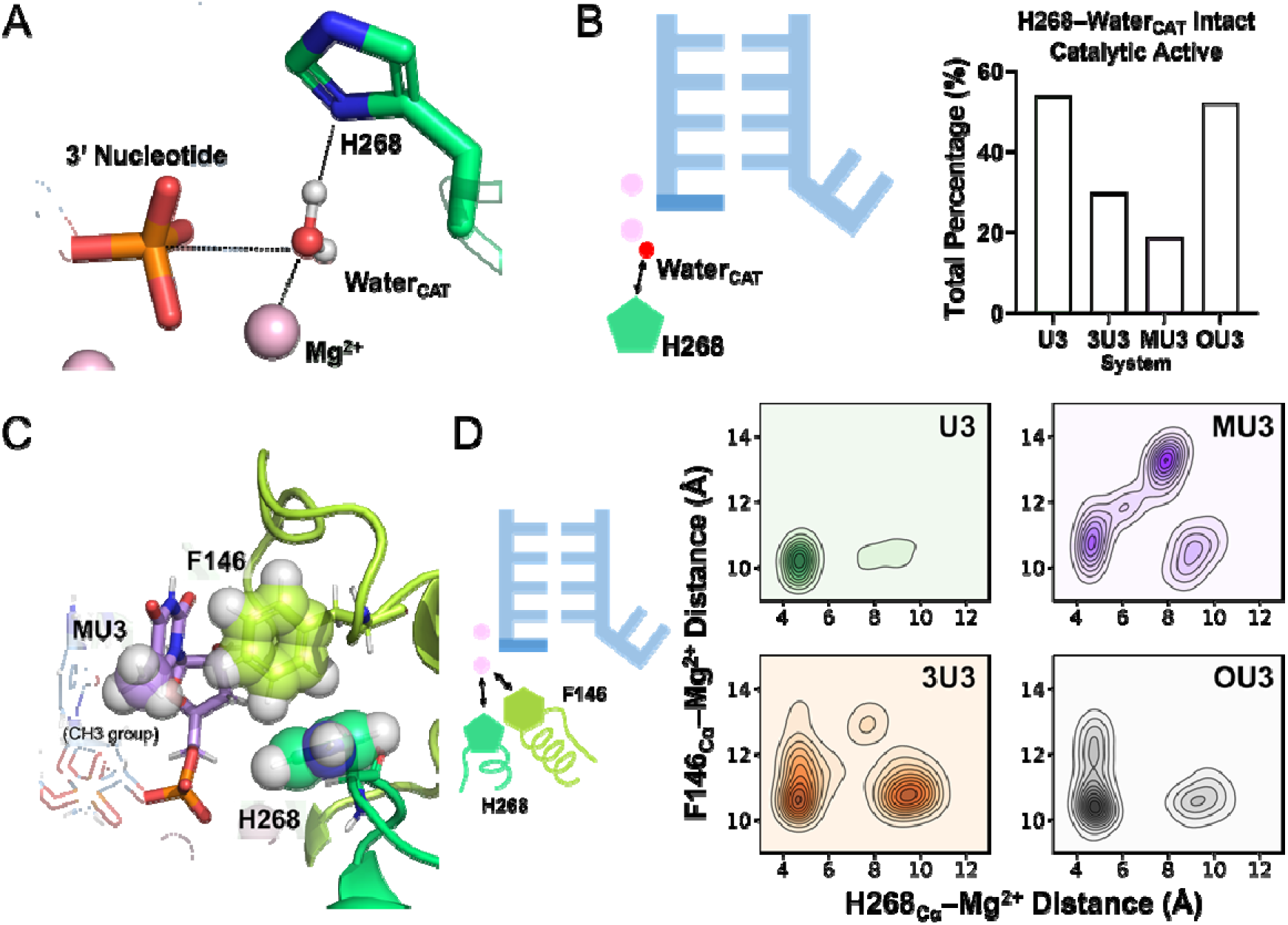
MD simulation on four systems of nsp14-nsp10-RNA complex containing terminal U, 3’-dU, 5-Methyl-3’-dU and 2’-O-Methyl-U. (A) Active site configuration of nsp14-nsp10-RNA complex in MD simulation model. (B) Total population of catalytic active configurations with criteria H268–Water_CAT_ distance < 3.5 Å. Water_CAT_ bound defined with criteria of Mg^2+^–Water_CAT_ distance < 2.2 Å and P3’Nucleotide–Water_CAT_ distance < 3.5 Å. (C) Close proximity between the 5-methyl group of 5-Methyl-3’-dU, F146 and H268 in active site. (D) 2D contour plot of H268_C_ –Mg^2+^ distance and F146 –Mg^2+^ distance. Abbreviation: U3=U, 3U3=3’-dU, MU3=5-Methyl-3’-dU, OU3=2’-O-Methyl-U.

The enhanced inhibition effect of 5-methylation could be attributed to the destabilization of F146 loop in the enclosed active site that is in close proximity with 3’ nucleotide and H268 (**Figure 8C**). Although being less recognized than other catalytic residues (e.g., D90, E92, or H268), F146 is also a conserved residue across coronaviruses^[43]^ and reported to be stacking with the nucleobase of 3’ nucleotide in SARS-CoV-2 nsp14-nsp10-RNA cryo-EM structure.^[41]^ From MD simulation, we discovered that 5-Methyl-3’-dU induced additional destabilization of F146 loop to 3’-dU. When using the stably bound catalytic Mg^2+^ as reference, the centering distance population of F146_C_α–Mg^2+^ shifted from ∼10 Å (U, 3’-dU and 2’-O-Methyl-U) to ∼13 Å (5-Methyl-3’-dU), with an associated opened H268 catalytic loop demonstrated by increased H268_C_α–Mg^2+^ distance (**Figure 8D**). Hence, the additional destabilization of F146 loop closure renders an extra barrier for active site closure in nsp14-nsp10-RNA complex with incorporated 5-Methyl-3’dU than 3’-dU. In total, our MD simulation suggested that nucleobase 5-methylation enhances resistance to exonuclease proofreading by destabilizing F146 stacking, which could serve a pan-CoV target for future drug development.

## 3. Discussion

Targeting the viral RdRp with nucleoside analogues remains a cornerstone strategy for antiviral development against coronaviruses, yet the nsp14-nsp10 proofreading machinery has critically limited the efficacy of this approach by excising incorporated nucleotide analogues from nascent viral RNA.^[22]^ In this study, we systematically evaluated nucleoside analogue scaffolds across multiple ribose and nucleobase modification sites, using a sequential two-tiered screening approach to identify compounds exhibiting both chain termination and proofreading resistance. Our findings identify 5-methyl-3′-dUTP as a standout inhibitor that satisfies both chain termination and proofreading resistance criteria, and provide mechanistic insight into the structural basis for its enhanced proofreading resistance.

Our systematic screening revealed distinct antiviral profiles associated with different modification sites. Among 2′-modified analogues, 2′-OMe-NTP functioned as an immediate chain terminator but was readily excised by nsp14-nsp10, indicating that 2′-O-methyl modification alone is insufficient to confer meaningful proofreading resistance. This observation is consistent with published data demonstrating that 2′-modifications generally provide limited protection against ExoN excision: while 2′-fluoro and 2′-C-methyl substitutions offer little to no resistance, and 2′-deoxy and 2′-amino provide only modest reductions in excision rates, all 2′-modifications tested exhibit substantially weaker ExoN resistance than 3′-deoxy modifications.^[44]^ In contrast, 3′-deoxy modifications consistently conferred both chain termination and meaningful ExoN resistance across all four canonical nucleobases tested, consistent with the established role of the 3′-OH in substrate recognition by the catalytic residue E92.^[41,44]^ Using an RdRp-directed incorporation system that directly recapitulates the physiological context of viral chain termination, our results systematically confirm this 3′-deoxy resistance mechanism across all four canonical nucleobases—an extent of coverage not previously reported. These findings directly address the suggestion that elimination of the 3′-OH may bolster anti-coronavirus nucleoside activity,^[44]^ and extend this concept by demonstrating that additional nucleobase modification at the 5-position can further augment both ExoN resistance and RdRp incorporation efficiency.

The distinct ExoN susceptibility profiles of 3′-dCMP and ddhCMP provide further insight into the structural basis of proofreading resistance. Although both analogues lack a canonical 3′-OH, published biochemical data demonstrate that 3′-dCMP-terminated RNA is substantially more resistant to nsp14-nsp10 cleavage than ddhCMP-terminated RNA, with the latter being excised nearly as efficiently as unmodified CMP.^[45]^ This divergence is likely explained by ribose conformation: whereas 3′-deoxyribose retains a flexible sugar pucker compatible with that of natural ribonucleotides, the ddhC ribose is constrained into a planar geometry by the C3′–C4′ double bond,^[44,45]^ which may preserve the structural complementarity required for productive engagement with the ExoN active site despite the absence of a 3′-OH. These observations reinforce the conclusion that 3′-deoxy modification confers genuine resistance to ExoN-mediated excision, and that this resistance depends critically on the overall three-dimensional geometry of the 3′-terminal ribose rather than simply the presence or absence of the hydroxyl group.

Within this framework, 5-methyl-3′-dUTP emerged as the standout candidate, demonstrating that nucleobase methylation can substantially augment the ExoN resistance already conferred by the 3′-deoxy scaffold. Direct comparison between 3′-dUMP and 5-methyl-3′-dUMP terminated RNAs revealed markedly superior resistance to nsp14-nsp10 cleavage for the latter, validated by both concentration-dependent and time-course analyses, fluorescence polarization measurements, and single-molecule FRET assays. Our MD simulations provide a mechanistic basis for this enhanced resistance. The loss of the 3′-OH reduced the population of catalytically competent configurations from 54% (unmodified U) to 30% (3′-dU), consistent with the established role of the 3′-OH in substrate recognition by E92.^[41]^ Strikingly, the 5-methyl group further reduced this population to 19% through an additional and independent mechanism: destabilization of the F146-containing loop in the nsp14-nsp10 active site, reflected by a shift in the F146_C_ –Mg^2+^ centering distance from ∼10 Å (U, 3′-dU, 2′-OMe-U) to ∼13 Å (5-methyl-3′-dU), with concomitant opening of the H268 catalytic loop. This two-pronged mechanism—disruption of the 3′-OH–E92 interaction compounded by F146 loop destabilization—collectively imposes a substantially higher barrier to active site closure relative to 3′-deoxy modification alone. These MD findings are consistent with, and provide dynamic mechanistic detail for, two recent structural studies that independently identified the same region of the ExoN active site as a key determinant of NA resistance. Wang *et al.* showed that insufficient interaction between the 3′-terminal nucleobase and the H95/Q145/F146 triad shifts the terminal nucleotides into a "huddle-up" conformation unfavorable for catalysis,^[46]^ while Liu *et al.* revealed at higher resolution that the 2′-CH_3_ group of incorporated NAs contacts P141 and F146 within the allosteric regulatory loop (P140–L149), displacing this loop and destabilizing the catalytically competent conformation of the ExoN active site.^[47]^ Although the two structural studies differ in their substrate conditions—Wang *et al.* used single-stranded RNA while Liu *et al.* employed the more physiologically relevant double-stranded RNA substrate, as noted by Liu *et al.*^[47]^—both converge on F146 and its surrounding loop as a critical structural element governing ExoN resistance, a conclusion consistent with our MD simulation data. Our results extend this framework by demonstrating that the 5-methyl group of a 3′-deoxy nucleotide is sufficient to destabilize this loop, providing a dynamic and quantitative basis for the design of ExoN-resistant nucleobase modifications. F146 is conserved across coronaviruses,^[47]^ suggesting that this resistance mechanism may be broadly applicable beyond SARS-CoV-2. Looking forward, the F146-containing allosteric loop could serve as a structural target for rational design of nucleobase modifications beyond 5-methylation, expanding the toolkit for developing proofreading-resistant antivirals across the coronavirus family.

In addition to its superior proofreading resistance, 5-methyl-3′-dUTP demonstrated substantially enhanced RdRp incorporation efficiency compared to its unmodified counterpart 3′-dUTP. In an incorporation assay measuring the apparent EC50 of RdRp-mediated incorporation in the absence of competing natural nucleotides, 5-methyl-3′-dUTP showed an approximately 7-fold improvement over 3′-dUTP (106.7 nM vs. 750.8 nM), indicating that the 5-methyl group substantially lowers the concentration required for effective RdRp utilization. Notably, the EC50 of 5-methyl-3′-dUTP (106.7 nM) is only approximately 2-fold higher than that of natural UTP, suggesting competitive incorporation even in the presence of the endogenous nucleotide pool. The mechanistic basis for this enhanced incorporation remains to be elucidated and will require structural studies of RdRp in complex with 5-methyl-3′-dUTP. The 5-methyl modification improves both proofreading resistance and RdRp incorporation efficiency. This convergence is rare among nucleoside scaffolds and represents a therapeutically valuable property.

Beyond proofreading resistance, the 3′-deoxy scaffold confers a second layer of antiviral durability: irreversibility of chain termination. Our cleavage-reextension assays demonstrated that RNA chains terminated with 3′-dUTP or 5-methyl-3′-dUTP could not be rescued even following nsp14-nsp10-mediated cleavage and supplementation with the correct nucleotide. This contrasts sharply with sofosbuvir-terminated RNA, which was efficiently rescued under identical conditions. The resistance to rescue of 3′-dUMP- and 5-methyl-3′-dUMP-terminated RNA stems primarily from their reduced susceptibility to nsp14-nsp10 cleavage. At moderate nsp14-nsp10 concentrations sufficient to cleave and rescue sofosbuvir-terminated RNA, both 3′-dUMP- and 5-methyl-3′-dUMP-terminated RNA showed minimal cleavage due to the inherent resistance of the 3′-deoxy termini, with the 5-methyl modification providing additional resistance to exonuclease activity, resulting in minimal re-extension. When nsp14-nsp10 concentrations were increased substantially to overcome this resistance, the cleaved 3′-deoxy-terminated RNA and 5-methyl-3′-deoxy-terminated RNA rapidly became vulnerable to extensive degradation by the high exonuclease activity. Under these conditions, the rate of RNA degradation exceeded the rate of RdRp-mediated re-extension, preventing productive rescue and maintaining the appearance of irreversible chain termination. The 3′-deoxy termini exhibit poor substrate properties for exonuclease cleavage, and the 5-methyl substitution further enhances this resistance. This creates a kinetic imbalance between exonuclease cleavage and RdRp re-extension: cleavage is either insufficient for rescue (at moderate nsp14-nsp10 concentrations) or so extensive that degradation outpaces re-extension (at high concentrations), preventing productive rescue in either scenario. This represents a fundamental advantage over chain-terminating analogues such as sofosbuvir, which are efficiently cleaved and rescued. The findings underscore the therapeutic rationale for prioritizing 3′-deoxy-based nucleoside analogues, particularly 5-methyl-3′-deoxy variants, which offer superior antiviral durability through enhanced resistance to exonuclease rescue.

In theory, the antiviral profile of 5-methyl-3′-dUTP compares favorably with those of existing approved therapeutics. Remdesivir acts as a delayed chain terminator and remains susceptible to nsp14-mediated excision; structural studies demonstrate that remdesivir incorporation destabilizes the RdRp–RNA interaction while concurrently enhancing RNA affinity for ExoN, thereby facilitating its removal from nascent viral RNA.^[22,48]^ In contrast, 5-methyl-3′-dUTP is an immediate chain terminator with markedly superior ExoN resistance, and its terminated RNA cannot be rescued even after partial cleavage. Molnupiravir exerts antiviral activity through lethal mutagenesis, which confers a high genetic barrier to resistance, but raises safety concerns: molnupiravir-induced mutational signatures have been identified in SARS-CoV-2 sequences from treated patients with evidence of onward transmission of drug-mutated viruses,^[49]^ and its active metabolite NHC can be incorporated into host mitochondrial RNA.^[50]^ Alternative chemical strategies, including α-phosphorothioate modification of the scissile phosphodiester bond, also confer complete ExoN resistance but at the cost of reduced incorporation efficiency and added complexity in prodrug design.^[51]^ The ribose-based mechanism described here—combining 3′-deoxy modification with 5-methyl nucleobase substitution—achieves ExoN resistance without modifying the phosphate backbone, and may therefore offer a more favorable metabolic and prodrug development profile.

This study has several limitations that define the roadmap for future work. Our experiments were conducted in a reconstituted in vitro system, and key parameters including intracellular phosphorylation efficiency, metabolic stability, cytotoxicity, and antiviral efficacy in cell culture remain to be established. As the active triphosphate form of 5-methyl-3′-dUTP cannot enter cells directly, the design of a cell-permeable prodrug represents the critical next step; phosphoramidate ProTide strategies, exemplified by the clinically approved antiviral remdesivir, provide a validated framework for intracellular delivery of nucleoside triphosphate analogues.^[52]^ While the potential for resistance mutations in RdRp or nsp14 was not investigated, prior clinical studies of remdesivir-treated cohorts consistently report that resistance mutations arise at low frequencies and are unlikely to contribute significantly to virologic rebound,^[53–58]^ providing reassurance for nucleoside-based therapeutic strategies more broadly.

Taken together, the dual mechanism of immediate chain termination and robust ExoN resistance identified here—augmented by enhanced RdRp incorporation efficiency and the irreversibility of the chain-terminated state—establishes 5-methyl-3′-dUTP as a structurally and mechanistically differentiated antiviral lead relative to currently approved therapeutics, and offers a starting point for the rational design of nucleoside analogues capable of circumventing the proofreading barrier that has broadly limited the efficacy of this therapeutic class against coronaviruses.

## 4. Conclusion

In conclusion, this study identifies 5-methyl-3′-dUTP as a promising antiviral lead compound distinguished by immediate chain termination, robust nsp14–nsp10-mediated proofreading resistance, and irreversibility of its chain-terminated state. MD simulations reveal that this resistance arises from a two-pronged mechanism involving disruption of the 3′-OH–E92 interaction and destabilization of the F146-containing allosteric regulatory loop, converging with independent structural studies^[46,47]^ to identify this conserved loop as a key pharmacological target for ExoN evasion across the coronavirus family. These findings offer a structural and mechanistic basis for the rational design of next-generation nucleoside analogues capable of circumventing coronavirus proofreading, with broader implications for the development of broad-spectrum antivirals as a durable component of pandemic preparedness.

## 5. Experimental Section/Methods

### Expression and Purification of the nsp12□nsp7□nsp8 Complex

The nsp12□nsp7□nsp8 complex was expressed and purified using the insect cell baculovirus system. A codon□optimised DNA sequence (BstEII□10×His tag nsp7□TEV□site□nsp8 TEV□site□nsp12□2×StrepII tag RsrII) was chemically synthesised and cloned into the pKL pBac backbone via BstEII and RsrII restriction sites, initiated by a 5′ ATG start codon and flanked by two Tobacco Etch Virus (TEV) protease cleavage sites for post translational processing. Recombinant baculovirus encoding the nsp12 nsp7 nsp8 complex was generated by transfecting the bacmid into High Five cells at a density of 2 × 10 cells mL ¹ cultured in ESF 921 serum free medium (Expression Systems) at 27 °C. After 5–7 days, the culture medium was clarified by centrifugation, filtered through a 0.22 µm filter, supplemented with 0.2% (w/v) bovine serum albumin (BSA), and stored at 4 °C protected from light as P0 virus stock. For protein expression, High Five cells at a density of 2–3 × 10 cells mL ¹ were infected with 5 mL of P0 virus in 200 mL culture volume and incubated for 3–5 days at 27 °C with shaking at 120 rpm. Cells were harvested by centrifugation at 500 × g for 5 min at 4 °C, and cell pellets were stored at −80 °C until use. For protein purification, cell pellets were resuspended in binding buffer containing 50 mM Tris HCl pH 7.4, 100 mM NaCl, 4 mM MgCl, 1 mM dithiothreitol (DTT), 10% (v/v) glycerol, and EDTA free Protease Inhibitor Cocktail (TargetMol). Cells were lysed by sonication on ice (BIOBASE Ultrasonic Cell Disruptor, 10 cycles of 15 s on/5 s off at 80% power rate), and the lysate was clarified by centrifugation at 50 000 rpm (Beckman Coulter Avanti JXN-26, JA-25.50 rotor) for 30 min at 10 °C. The supernatant was further clarified by filtration through a 0.45 µm syringe filter. The clarified lysate was loaded onto a gravity flow column packed with Strep Tactin®XT 4Flow® resin (IBA Lifescience) pre equilibrated with 10 column volumes (CV) of wash buffer (50 mM Tris HCl pH 7.4, 100 mM NaCl, 4 mM MgCl, 1 mM DTT, and 10% (v/v) glycerol). After sample loading, the column was washed with 10 CV of wash buffer, and bound protein was eluted with elution buffer (50 mM Tris HCl pH 7.4, 100 mM NaCl, 4 mM MgCl, 1 mM DTT, 100 mM biotin and 2 mM EDTA). Fractions containing the nsp12 nsp7 nsp8 complex were identified by SDS PAGE analysis. The pooled Strep Tactin eluate was diluted twofold with low salt buffer (50 mM Tris HCl pH 7.4, 50 mM NaCl, 4 mM MgCl, 1 mM DTT) and loaded onto a HiTrap Heparin HP affinity column (1 mL, Cytiva) pre equilibrated in low salt buffer using an ÄKTA FPLC system. The column was washed with 5 CV of low salt buffer, and the nsp12 nsp7 nsp8 complex was eluted using a linear gradient from 50 mM to 1 M NaCl over 30 CV at a flow rate of 1 mL min ¹. Peak fractions were analysed by SDS PAGE. The pooled protein was concentrated and buffer exchanged into storage buffer (50 mM Tris HCl pH 7.4, 150 mM NaCl, 1 mM DTT) by repeated cycles of concentration and dilution using Amicon Ultra 15 centrifugal filter units (30 kDa MWCO, Millipore). Protein concentration was determined using the Pierce™ Dilution-Free™ Rapid Gold BCA Protein Assay (Thermo Fisher Scientific) with bovine serum albumin as a standard. The final protein sample was aliquoted, flash frozen in liquid nitrogen, and stored at −80 °C.

### Expression and Purification of the nsp14 nsp10 Complex

The nsp14 nsp10 complex was expressed and purified using the insect cell baculovirus system. A codon optimised DNA sequence (BstEII 10×His tag nsp14 TEV site nsp10 2×StrepII tag RsrII) was chemically synthesised and cloned into the pKL pBac backbone via BstEII and RsrII restriction sites. The expression and purification procedures were conducted as described for the nsp12 nsp7 nsp8 complex with minor modifications as detailed below. Recombinant baculovirus encoding the nsp14 nsp10 complex was generated by transfecting the bacmid into High Five cells at a density of 2 × 10 cells mL ¹ cultured in ESF 921 serum free medium (Expression Systems) at 27 °C. After 5–7 days, the culture medium was clarified by centrifugation, filtered through a 0.22 µm filter, supplemented with 0.2% (w/v) bovine serum albumin (BSA), and stored at 4 °C protected from light as P0 virus stock. For protein expression, High Five cells at a density of 2–3 × 10 cells mL ¹ were infected with 5 mL of P0 virus in 200 mL culture volume and incubated for 3–5 days at 27 °C with shaking at 120 rpm. Cells were harvested by centrifugation at 500 × g for 5 min at 4 °C, and cell pellets were stored at −80 °C until use. For protein purification, cell pellets were resuspended in binding buffer containing 50 mM Tris HCl pH 7.4, 100 mM NaCl, 5 mM MgCl, 10% (v/v) glycerol, and EDTA free Protease Inhibitor Cocktail (TargetMol). Cells were lysed by sonication on ice (BIOBASE Ultrasonic Cell Disruptor, 10 cycles of 15 s on/5 s off at 80% power rate), and the lysate was clarified by centrifugation at 50 000 rpm (Beckman Coulter Avanti JXN-26, JA-25.50 rotor) for 30 min at 10 °C. The supernatant was further clarified by filtration through a 0.45 µm syringe filter. The clarified lysate was loaded onto a gravity flow column packed with BeyoGold His tag Purification Resin (Beyotime) pre equilibrated with 10 column volumes (CV) of wash buffer (50 mM Tris HCl pH 7.4, 100 mM NaCl, 5 mM MgCl, 10% (v/v) glycerol). After sample loading, the column was washed with 10 CV of wash buffer, and bound protein was eluted with elution buffer (50 mM Tris HCl pH 7.4, 100 mM NaCl, 5 mM MgCl, 250 mM imidazole). Fractions containing the nsp14 nsp10 complex were identified by SDS PAGE analysis. The pooled Ni NTA eluate was directly loaded onto a gravity flow column packed with Strep Tactin®XT 4Flow® resin (IBA Lifescience) pre equilibrated with 10 CV of wash buffer (50 mM Tris HCl pH 7.4, 100 mM NaCl, 5 mM MgCl, 1 mM DTT, 10% (v/v) glycerol). After sample loading, the column was washed with 10 CV of wash buffer, and bound protein was eluted with elution buffer (50 mM Tris HCl pH 7.4, 100 mM NaCl, 5 mM MgCl, 1 mM DTT, 50 mM biotin, and 2 mM EDTA). The pooled protein was concentrated and buffer exchanged into storage buffer (50 mM Tris HCl pH 7.4, 150 mM NaCl, 1 mM DTT) by repeated cycles of concentration and dilution using Amicon Ultra 15 centrifugal filter units (30 kDa MWCO, Millipore). Protein concentration was determined using the Pierce™ Dilution-Free™ Rapid Gold BCA Protein Assay (Thermo Fisher Scientific) with bovine serum albumin as a standard. The final protein sample was aliquoted, flash frozen in liquid nitrogen, and stored at −80 °C.

### RdRp Enzymatic Activity Assay and Inhibition by Nucleotide Analogues

Primer extension assays were performed using a fluorescently labeled 20 nucleotide RNA oligonucleotide (LS2U, 5′ FAM GUCAUUCUCCUAAGAAGCUU 3′) as the primer strand and two different template strands: a 40 nucleotide RNA oligonucleotide LS15A (5′ CUAUCCCCAUGUGAUUUUACAAGCUUCUUAGGAGAAUGAC 3′) for testing GTP and UTP analogues, and a 41 nucleotide RNA oligonucleotide LS41 GU (5′ CAGUAAGAGGAUUCUUCGAAGUCCUUUAGUGUACCCCUAUC 3′) for testing CTP and ATP analogues. RNA oligonucleotides were purchased from Tsingke Biotech and resuspended in RNase free water. The primer and template were annealed at a 1:1 molar ratio by heating to 75 °C for 5 min followed by slow cooling to room temperature over 10 min. Polymerase reactions were performed in a final volume of 10 µL containing 1 µM purified nsp12 nsp7 nsp8 complex, 0.1 µM annealed primer template duplex, and 25 µM of each ribonucleoside triphosphate (rNTP) in reaction buffer (20 mM Tris HCl pH 7.5, 5 mM MgCl). For inhibition assays, individual nucleotide analogues were added at a final concentration of 100 µM, replacing the corresponding natural NTP. All reactions were prepared using RNase free water and incubated at 37 °C for 10 min. Reactions were quenched by adding an equal volume (10 µL) of 2× RNA loading buffer containing 95% (v/v) formamide, 18 mM EDTA, and 0.025% (w/v) each of xylene cyanol and bromophenol blue. Samples were heated at 95 °C for 3 min and immediately placed on ice. A total of 8 µL of each sample was loaded onto a 15% denaturing polyacrylamide gel containing 8 M urea in 1× TB buffer (130 mM Tris, 45 mM boric acid). Electrophoresis was performed in 0.5 × TB buffer at 350 V for 60 min at room temperature. Gels were imaged using a LI COR Odyssey M Imager with detection at 488 nm (FAM channel).

### Exonuclease Activity Assay

To assess the susceptibility of nucleotide analogue terminated RNA to exonuclease mediated excision, a two step assay was performed. First, RNA substrates terminated with specific nucleotide analogues were generated by primer extension reactions. The 5′ fluorescein labeled primer (LS2U, 0.1 µM) was annealed with the template strand (LS15A, 0.1 µM) as described in the “RdRp Enzymatic Activity Assay and Inhibition by Nucleotide Analogues” section. Polymerase reactions were carried out in a final volume of 10 µL containing 1 µM purified nsp12 nsp7 nsp8 complex in reaction buffer (20 mM Tris HCl pH 7.5, 5 mM MgCl). For reactions with natural nucleotides, all four rNTPs were added at 25 µM each. For analogue incorporation, the corresponding natural NTP was replaced with the respective nucleotide analogue at 100 µM, while the other three natural rNTPs remained at 25 µM. Reactions were incubated at 37 °C for 10 min to generate analogue terminated RNA products. Following the polymerase reaction, exonuclease activity was assessed by adding the purified nsp14 nsp10 complex directly to the reaction mixture without prior purification of the RNA products. For concentration dependent experiments, varying concentrations of nsp14 nsp10 complex (ranging from 0 to 300 nM) were added, and the reactions were incubated at 37 °C for 10 min. For time course experiments, a fixed concentration of 150 nM nsp14 nsp10 complex was added, and aliquots were collected at various time points (0, 5, 10, 20, and 40 min) during incubation at 25 °C. All reactions were quenched by adding an equal volume (10 µL) of 2× RNA loading buffer containing 95% (v/v) formamide, 18 mM EDTA, and 0.025% (w/v) each of xylene cyanol and bromophenol blue. Samples were heated at 95 °C for 3 min and immediately placed on ice. A total of 8 µL of each sample was loaded onto a 15% denaturing polyacrylamide gel containing 8 M urea in 1× TB buffer (130 mM Tris, 45 mM boric acid). Electrophoresis was performed in 0.5 × TB buffer at 350 V for 30 min at room temperature. Gels were imaged using a LI COR Odyssey M Imager with detection at 488 nm (FAM channel). Band intensities were quantified using Image Lab software (Bio Rad Laboratories). The extent of exonuclease activity was determined by measuring the percentage of remaining uncleaved RNA substrate.

### Exonuclease Activity Assay by Fluorescence Polarisation

The exonuclease activity of the SARS CoV 2 nsp14 nsp10 complex and its sensitivity to nucleotide analogues were assessed in real time using a fluorescence polarisation (FP) based assay. The fluorophore labeled RNA substrates, FAM LS2U G 3′ dU and FAM LS2U G 5 Methyl 3′ dU, were enzymatically synthesised by the SARS CoV 2 RNA dependent RNA polymerase (RdRp). Briefly, reactions contained 20 mM Tris HCl pH 7.5, 5 mM MgCl, 0.1 µM FAM LS2U primer, 0.1 µM LS15A template, 1.5 µM nsp12 nsp7 nsp8 complex, 25 µM GTP, and 25 µM UTP or its respective analogue (3′ dUTP or 5 Methyl 3′ dUTP). After incubation at 37 °C for 10 min, the products were purified using a Monarch® Spin RNA Cleanup Kit (10 μg) (New England Biolabs, T2030L) and eluted in nuclease free water. The concentration of the purified substrates was determined using a DeNovix Spectrophotometer / Fluorometer (DS-11 FX+), and their integrity was confirmed by denaturing PAGE. Real time exonuclease reactions were performed in black, round bottom 384 well plates (Corning) in a total reaction volume of 5 µL. The standard reaction buffer consisted of 20 mM Tris HCl pH 7.5, 5 mM MgCl, and 5% (v/v) glycerol. Each well contained 0.05 µM of the specified FAM labeled RNA substrate. Reactions were initiated by the addition of 10 nM nsp14 nsp10 complex. The plate was immediately transferred to a pre equilibrated BMG LABTECH CLARIOstar microplate reader maintained at 25 °C. Fluorescence polarisation (measured in millipolarisation units, mP) was monitored every minute for a period of approximately 20 min. The raw mP values were exported and analysed in GraphPad Prism (version 10.0).

### Nucleotide Analogue Rescue Assay

To assess whether nucleotide analogue terminated RNA products could be rescued by the exonuclease proofreading activity and subsequently extended to full length products, a multi step rescue assay was performed with modifications based on previously published methods.^[39]^ This assay evaluates the combined action of nsp14 nsp10 mediated excision and nsp12 nsp7 nsp8 mediated re extension in the presence of natural nucleotides. In the first step, primer extension reactions were performed to generate analogue terminated RNA products. Reactions were carried out in a final volume of 10 µL containing 1.5 µM purified nsp12 nsp7 nsp8 complex, 0.1 µM annealed FAM LS2U/LS15A primer template duplex, 4 µM each of ATP, CTP, and GTP, and 25 µM of the nucleotide analogue (sofosbuvir TP, 3′ dUTP, or 5 methyl 3′ dUTP) in reaction buffer (20 mM Tris HCl pH 7.5, 5 mM MgCl). Reactions were incubated at 37 °C for 20 min to allow incorporation of the nucleotide analogues and generation of stalled RNA products. Following the initial polymerase reaction, the rescue step was initiated by simultaneously adding varying concentrations of nsp14 nsp10 complex (ranging from 0 to 150 nM) and 4 µM natural UTP to the reaction mixture without prior purification or buffer exchange. The reactions were incubated for an additional 30 min at 37 °C to allow exonuclease mediated excision of the terminal nucleotide analogue and subsequent re extension by the polymerase in the presence of natural UTP. Control reactions without nsp14 nsp10 addition or without natural UTP supplementation were performed in parallel to distinguish between direct extension and rescue mediated extension. Reactions were quenched by adding an equal volume (10 µL) of 2× RNA loading buffer containing 95% (v/v) formamide, 18 mM EDTA, and 0.025% (w/v) each of xylene cyanol and bromophenol blue. Samples were heated at 95 °C for 3 min and immediately placed on ice. A total of 8 µL of each sample was loaded onto a 15% denaturing polyacrylamide gel containing 8 M urea in 1× TB buffer (130 mM Tris, 45 mM boric acid). Electrophoresis was performed in 0.5 × TB buffer at 350 V for 30 min at room temperature. Gels were imaged using a LI COR Odyssey M Imager with detection at 488 nm (FAM channel). Band intensities were quantified using Image Lab software (Bio Rad Laboratories). The extent of rescue was determined by comparing the ratio of full length extension products in the presence versus absence of nsp14 nsp10.

### RNA for single-molecule FRET assay

We ordered RNA oligonucleotides from Integrated DNA Technologies (IDT), conjugated to the fluorescent labels Cy3 or Cy5. The sequence of the original template used was 5’ - /Cy3/AAAAAAAAUUUUUUUUUUUUUUUAAUUCUUAAUCUCACAUAGC - 3’, while the sequence of the modified template was 5’ - /Cy3/UUUUUUUUAAAAAAAAAAAAAAAUUCUUAAUCUCACAUAGC - 3’. The sequence of the primer was 5’ - /Cy5/GCUAUGUGAGAUUAAGAAUU - 3’. Template and primer RNAs were annealed at a final concentration of 300 nM in hybridization buffer [20 mM Tris-HCl pH 8.0, 1 mM ethylenediaminetetraacetic acid (EDTA), 500 mM NaCl] using a single cycle temperature gradient from 95°C to 4°C.

### Protein for single-molecule assays

The expression and purification of nsp7, nsp8 and nsp12 for single-molecule assays has been described previously.^[40]^ The SARS-CoV-2 nsp10-nsp14-His6 construct was cloned into a MultiBac baculovirus with the pFL transfer vector.^[59]^ Colonies positive for both nsp10 and nsp14 by colony PCR were mini prepped using the NEB mini-prep kit before precipitating the DNA 1:1 with propan-2-ol at -20 °C. The DNA was then washed with ethanol twice and the pellet left to dry before resuspending in 100 μL Sf9 media (Gibco) and 20 μL lipofectin (Thermo). Sf9 cells were seeded at 0.5 million cells/mL in a 6-well plate and left for 30 mins to ensure adherence. The Sf9 media was removed and replaced by 60 μL of the resuspended pellet followed by 3 mL Sf9 media. The plate was sealed with parafilm and incubated at 27 °C for 62 h. 3mL baculovirus V1 was collected from the supernatant and used to infect 50 mL of Sf9 cells at 0.5 mil/mL. The cells were maintained at 0.5 mil/mL in order to reach 1 mil/mL at the end of a 4-day period for collection of 50 mL of baculovirus V2. Expansion cultures were made by seeding Sf9 cells at 0.005mil/mL overnight, then infecting with 2.5 mL baculovirus V2 per 500 mL culture. After 3 days, the culture was centrifuged using the JA 8.1 Rotor at 2500 x g for 15 mins at 4 °C.

The pellet was resuspended in wash buffer [50 mM HEPES-NaOH, pH 7.5, 300 mM NaCl, 10% glycerol, 0.05% n-Octyl β-d-thioglucopyranoside (OTG), 1 mM dithiothreitol (DTT)] with 1 pill of protease inhibitor (SIGMAFAST) and 1:1000 RNase A] before cell lysis by sonication at 40W pulse power, with pulsing for 50% of the 4 min duration. The cell lysate was centrifuged at 35 000 × g for 45 mins at 4°C and the supernatant was incubated with 1 ml Ni-NTA agarose (Merck Life Sciences) in wash buffer per litre of cell culture for 3–4 h at 4°C. The solution was washed three times with Imidazole Wash Buffer [50 mM HEPES-NaOH, pH 7.5, 400 mM NaCl, 20 mM Imidazole, 5% glycerol, 1 mM dithiothreitol (DTT)] by spinning at 2000 rpm for 4 mins and resuspending the beads. The his-tagged protein was then eluted from the beads in Imidazole Elution Buffer [50 mM HEPES-NaOH, pH 7.5, 400 mM NaCl, 90 mM Imidazole, 5% glycerol, 1 mM dithiothreitol (DTT)] for 10 mins at 4°C and then a final spin at 2000 rpm for 4 mins to isolate the supernatant. The imidazole was removed by dialysis in Dialysis Buffer [50 mM HEPES-NaOH, pH 7.5, 120 mM NaCl, 5% Glycerol, 1 mM dithiothreitol (DTT)]. The next day, the supernatant was concentrated to 0.5–2 ml and loaded on to a Superdex 200 10/300 increase column in size exclusion chromatographic (SEC) running buffer, nsp10-14 SEC Buffer [25 mM HEPES-NaOH, pH 7.5, 120 mM NaCl, 5% glycerol, 1 mM dithiothreitol (DTT)]. SEC fractions were checked by sodium dodecyl sulfate–polyacrylamide gel electrophoresis (SDS–PAGE) against the pre-stained precision plus 10–250 kDa Bio-Rad ladder (161–0373) and those with target protein were collected and concentrated to ∼5–10 mg/ml with an Amicon Millipore MWCO tube (10 kDa cutoff), aliquoted and flash-frozen in liquid nitrogen before storage at −80°C.

### Single-molecule fluorescence spectroscopy

The EI-FLEX confocal microscope (Exciting Instruments) was used for single-molecule FRET experiments using alternating-laser excitation (ALEX) to excite donor and acceptor fluorophores sequentially. Final concentrations of 8.5 µM RdRp were incubated with 10 nM pre-annealed RNA for 30 min at 30°C in a 3 μl reaction containing master mix (MM) (5 mM MnCl2, 300 µM ATP, 300 µM UTP, 10 mM KCl, 1 U/μl RNasin, 0.1 mM DTT) and protein storage buffer (PSB) (6.25 mM HEPES, 37.5 mM NaCl, 0.5 mM MnCl2, 0.25 mM DTT, 20% glycerol). For low NTP concentration reactions, a final concentration of 10 µM ATP and was 10 µM UTP used. The RdRp/RNA complex was then diluted to a final RNA concentration of ∼50 pM in confocal buffer (MnCl2 7 mM, KCl 10 mM, RNAsin 0.4 U/μl, DTT 0.2 mM, HEPES, pH 7.5 25 mM, NaCl 100 mM in water) for confocal imaging and analysis. For the nsp14-nsp10 assay, a volume of 6 μl reaction mixture containing RdRp, RNA, and MM containing 10 µM ATP/UTP was produced as above. After 20 minutes of incubation at 30°C, 3 μl was removed and imaged. The additional 3 μl of reaction mixture was combined with 3 μl of nsp14-nsp10 at a final concentration of 4 μM and further master mix. The secondary mixture was incubated for a further 20 minutes at 30°C before imaging. 3’-dUTP and 5-methyl-3’-dUTP were initially diluted into PSB prior to addition to master mixes when used, at the concentrations indicated in the figure legends.

We used a 2-laser line set up with 30–40 mW lasers at 520 and 638 nm. The 520 nm laser was used at 0.22 mW power with a 100 μs alternation period: OFF: 0 μs, ON: 45 μs, OFF: 55 μs, while the 638 nm laser power was 0.16 mW and alternating OFF: 50 μs, ON: 45 μs, OFF: 5 μs. The Exciting Instruments software was used to measure and evaluate the signals obtained.

### Artificial Intelligence Tools

During the preparation of this manuscript, AI writing assistance tools were used by some of the authors. Specifically, Claude (claude-sonnet, Anthropic) was used to improve the language and readability of the text. All AI-assisted content was reviewed and edited as needed. The authors take full responsibility for the content of the publication.

## Author Contributions

Peter Pak-Hang Cheung conceived the study. Lin Yang performed gel-based and fluorescence polarization assays and data analysis. Rory Cunnison performed smFRET assays and data analysis. Carmen Ka Man Tse and Tiantian Xu performed MD simulations and analysis. Buyu Zhang, Rick Xinzhou Xu, Daqi Yu, and Mingyuan Li assisted with experimental design and troubleshooting. Danielle Groves, Benny Zhibin Liang, Xunyu Zhou, Adrian Deng, and Jeremy R. Keown assisted with experiments. Lin Yang, Rory Cunnison, and Carmen Ka Man Tse drafted the original manuscript. Carmen Ka Man Tse, Daqi Yu, Nicole C. Robb, Yuanliang Zhai, and Billy Wai-Lung Ng reviewed and revised the manuscript. Peter Pak-Hang Cheung, Nicole C. Robb, and Lu Zhang supervised the project and secured funding. All authors read and approved the final manuscript.

## Acknowledgements

We thank the Li Ka Shing Translational Omics Platform for technical support. This work was supported by Collaborative Research Fund (C4032-21G) and Early Career Scheme (24303324) for P. P. C., National Natural Science Foundation of China/Research Grants Council of Hong Kong Joint Research Scheme (N_CUHK484/23) for P. P. C. and X. X. The Hong Kong RGC Postdoctoral Fellowship Scheme (PDFS2324-4S06) for X. X. We thank Professor Ervin Fodor and Dr Haitian Fan from the University of Oxford for the kind gift of the nsp10-nsp14-His6 plasmid.

## Data Availability Statement

The data that support the findings of this study are available from the corresponding author upon reasonable request. Received: ((will be filled in by the editorial staff)) Revised: ((will be filled in by the editorial staff)) Published online: ((will be filled in by the editorial staff))

## Supporting Information

Supporting Information is available from the Wiley Online Library or from the author.

