## Supporting Information for "Nucleobase Methylation Enhances SARS-CoV-2 Chain Terminator Evasion of Exonuclease Proofreading"

* Corresponding authors.

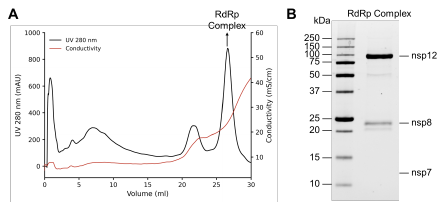

**Figure S1.** Purification of the SARS-CoV-2 nsp12-nsp7-nsp8 complex. (A) Heparin affinity chromatography elution profile. UV absorbance at 280 nm (black) and conductivity (red) are plotted against elution volume. The major peak at ~27 ml (arrow) corresponds to the eluted nsp12-nsp7-nsp8 complex. (B) SDS-PAGE analysis of the peak fraction. Bands Corresponding to nsp12 and nsp8 are visible; nsp7 is not apparent on the gel.

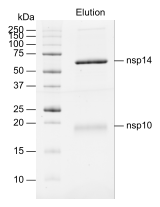

**Figure S2.** Purification and characterization of the nsp14-nsp10 complex. SDS-PAGE analysis of the purified nsp14-nsp10 complex. The elution fraction shows two distinct bands corresponding to nsp14 (~70 kDa) and nsp10 (~20 kDa), confirming successful co-purification of the complex.

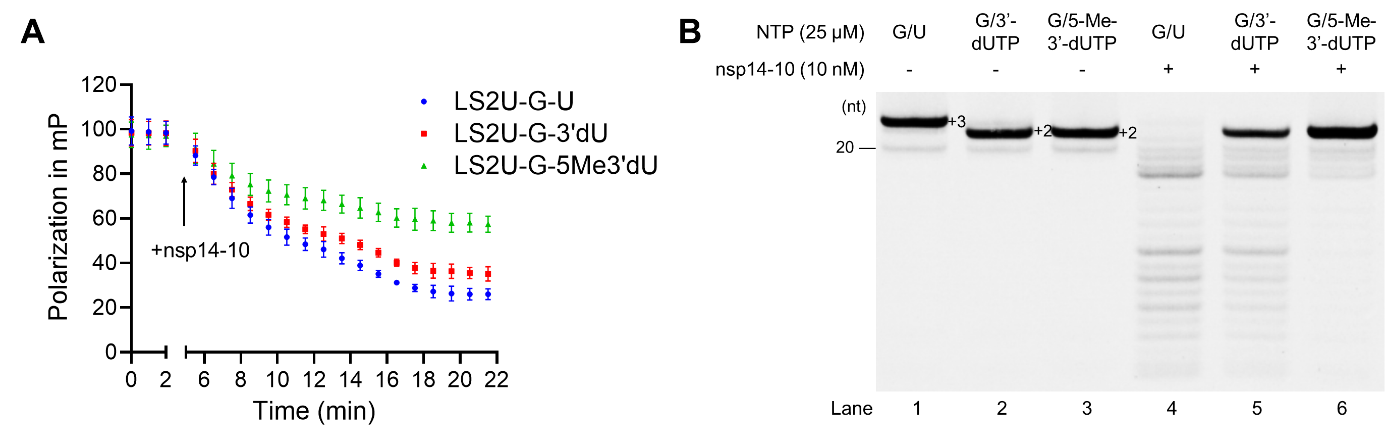

**Figure S3.** Real-time analysis of nsp14-nsp10 exonuclease activity by fluorescence polarization. (A) Real-time fluorescence polarization (FP) analysis of nsp14-nsp10 cleavage. Real-time FP signals were measured for fluorophore-labeled RNA substrates containing UMP (natural residue), 3'-dUMP, or 5-methyl-3'-dUMP residuals upon the addition of the nsp14-nsp10 complex. The differential decrease in FP signal reflects the relative resistance of the substrates to exonucleolytic cleavage. (B) Endpoint analysis of reaction products by denaturing PAGE. Denaturing polyacrylamide gel electrophoresis (PAGE) analysis of the endpoint reaction products from the real-time FP assays shown in panel (A). This confirms the relative final cleavage efficiency of the substrates. Data are presented as means ± SD from three independent experiments (n = 3).

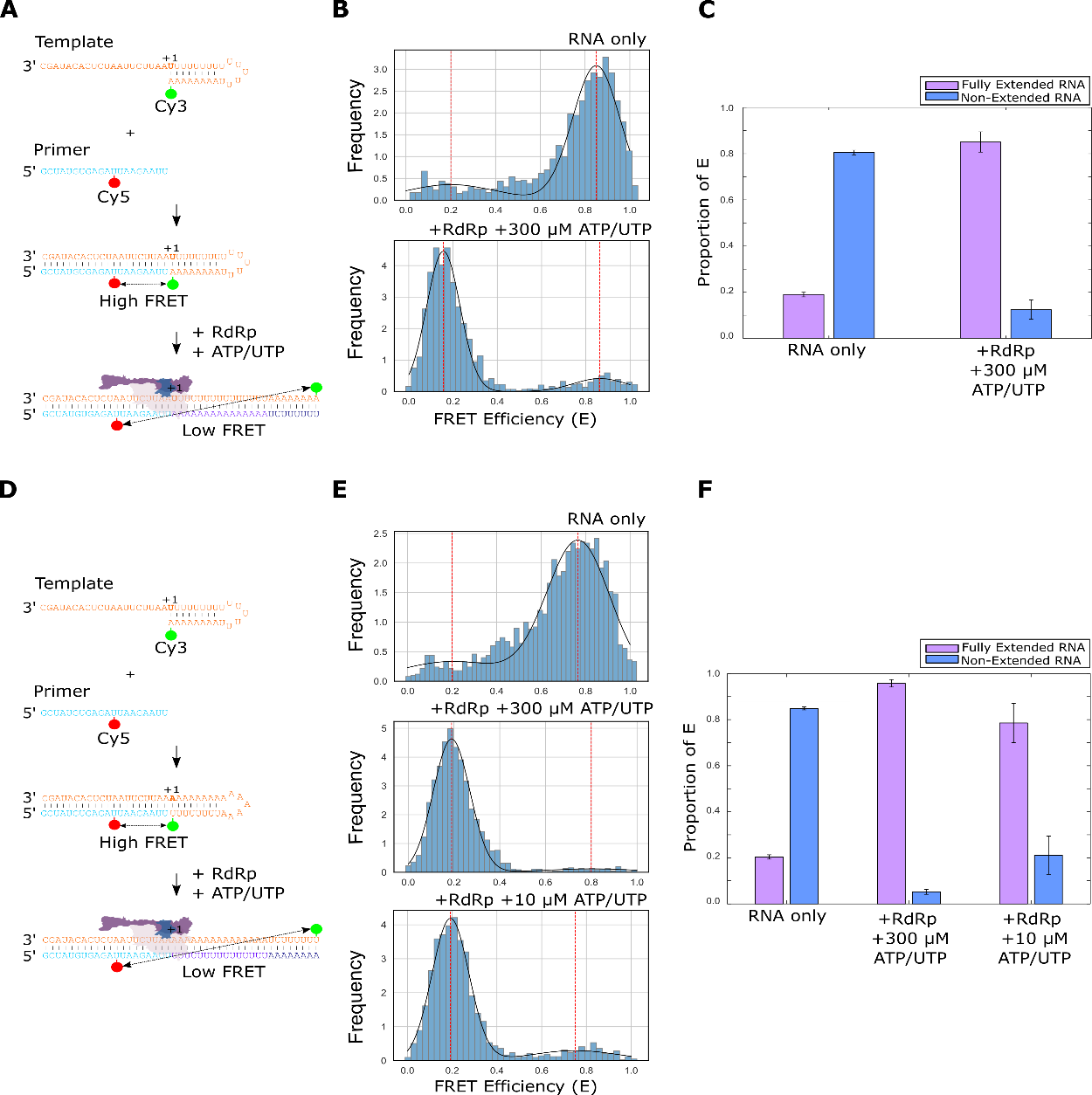

**Figure S4.** Design of a single-molecule FRET assay to detect RNA extension. (A) Schematic showing the original design of a single-molecule extension assay.^[1]^ A 43-nucleotide extension RNA template designed to form a self-annealed hairpin labelled with a Cy3 dye at the 5' end was annealed to a short 20-nucleotide primer sequence, labelled with a Cy5 dye at position 12 (high FRET signature). Upon RdRp and nucleotide addition the primer sequence is extended, separating the two dyes (low FRET signature). (B) FRET efficiency (E) when a Cy3-labelled 43mer template and Cy5-labelled 20mer primer were annealed without RdRp (top) or with RdRp and 300 µM ATP/UTP (bottom). (C) Quantification of the high and low FRET populations in panel (B) as a proportion of the total FRET distributions by finding the probability density function. Error bars represent standard error of the Gaussian fit. (D) Schematic of RNAs used to test the specific addition of UTP, with a modified template sequence so that UTP or its analogues are incorporated first. (E) Confirmation that the modified template works in a similar manner to the original assay design, with both 300 and 10 µM ATP/UTP. (F) Quantification of the high and low FRET populations in panel (E).

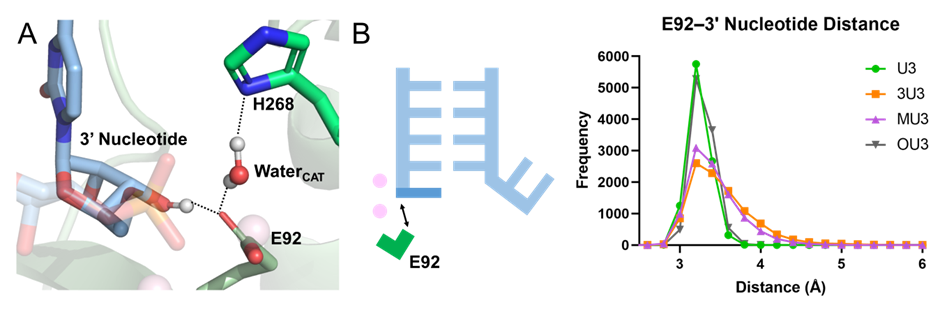
**Figure S5.** Effect of 3’-deoxy modification in enclosed active site in MD simulations. (A) Active site configuration of nsp14/10-RNA complex in MD simulation model. (B) Distance between the C3’ atom of terminal nucleotide and E92 in the enclosed active site.

**1. Structural modelling**

The cryo-EM structure of SARS-CoV-2 nsp14/10 (E191A)-RNA complex (PDBID: 7N0C) with Mg^2+^ ions was used to construct the model system for MD simulation. Side chain coordinates of E191 were taken from another complex (PDBID: 7N0B^[2]^) to restore as WT. The missing loop of nsp14 was modelled by structural alignment with a cryo-EM structure (PDBID: 7EIZ^[3]^) containing a complete loop (resid 446 to 489) and the coordinating Zn^2+^ ion, with the missing side chain of H455 modelled with Coot 0.9.8.^[4]^ The protonation states of histidine residues were assigned using PDB2PQR v3.6.1.^[5]^ with manual inspection to maintain coordination with the structural Zn^2+^ ions. Protonation state of cysteine residues coordinating with Zn^2+^ ions were also preassigned. The template and product RNA molecules were truncated to 15-nt and 14-nt respectively to reduce system size.

**2. Molecular dynamics simulation**

**2.1. Force field parameters** Amber ff14SB protein force field,^[6]^ OL15 for DNA^[7–9]^ and χOL3 for RNA^[10,11]^ were used. For 3’-dU, 5-Methyl-3’dU and 2'OMe-U, force field parameters were derived to be compatible with the chosen force field. In particular, a common set of RESP charges for modified ribose was derived by cross-fitting with A/U/C/G nucleotide molecules following a similar scheme as used in the original paper^[12]^ for generality. This approach was used for both 3’-dN and 2’OMe-N. The initial configuration used for nucleotides in force field parameters development were built from high resolution x-ray structures containing respective natural NTPs, including: ATP (PDBID: 4FF3^[13]^), UTP (PDBID: 5VZB^[14]^), CTP (PDBID: 4DQQ^[15]^) and GTP (PDBID: 6UOL^[16]^). Geometry optimization and single-point calculations were performed at HF/6-31G* level of theory with GAMESS 2023 R2.^[17]^ For both 3’-dN and 2’OMe-N, RESP charges equivalence was enforced across the four nucleotides on the ribose atoms excluding C1’ and H1’. To maintain compatibility with current Amber force field, H5T, O5’, O3’ and H3T atoms were assigned as 0.42950, -0.62230, -0.65410 and 0.43760 respectively. Considering 3’-dN lacks O3’–H3T group for extension, its 3’ end group was not enforced with pre-assigned value due to its inability to connect to the next nucleotide. RESP charges for 5-Methyl-3’dU were derived based on 3’dN. The partial charges of ribose (excluding C1’ and H1’) were enforced as those from the universal 3’dN set, while those of nucleobase fitted according to the default RESP scheme. All bonded parameters were taken from existing Amber force field, with nucleobase parameters of 5-Methyl-3’dU taken from T.

**2.2. MD simulation** Four representative 3’-end nucleotides were studied, including: (1) natural U (control), (2) 3’-dU, (3) 5-Methyl-3’dU and (4) 2'OMe-U (negative control). The original model containing 3’-end natural C (from section 1) was first equilibrated and used to generate universal initial structure for all the model systems. This model was solvated in dodecahedron box with 1.2 nm boundary. Neutralizing Mg^2+^ and Cl^-^ ions were added at concentration of 0.15 M. Harmonic restraint (1×10^5^ kJ mol^-1^ nm^-2^) was added between structural Zn^2+^ ions and coordinating residues to maintain coordination, with *b_0_*(NB–Zn)=0.21 nm and *b_0_*(S–Zn)=0.23 nm adopted from high resolution crystal structures.^[18]^ The system was gradually relaxed and fully equilibrated as followings:

(i)	10,000 steps of energy minimization using steepest decent with position restraint (1×10^3^ kJ mol^-1^ nm^-2^) on all the heavy atoms of protein/RNA and the structural Zn^2+^ ions, and restraint (1×10^4^ kJ mol^-1^ nm^-2^) on the two catalytic Mg^2+^ ions;

(ii)	10,000 steps of energy minimization using steepest decent with position restraint (1×10^4^ kJ mol^-1^ nm^-2^) on the two catalytic Mg^2+^ ions;

(iii)	200 ps of NVT equilibration (T=300 K) with position restraint (1×10^3^ kJ mol^-1^ nm^-2^) on all the heavy atoms of protein/RNA, the structural Zn^2+^ ions and the two catalytic Mg^2+^ ions, with additional harmonic restraint (1×10^5^ kJ mol^-1^ nm^-2^) between catalytic Mg^2+^ ions and coordinating residues (*b_0_* taken from (ii)) to reduce perturbation on Mg^2+^ during relaxation;

(iv)	500 ps of NPT equilibration (T=300 K, P=1 bar) with the same restraints as (iii).

(v)	20 ns of NPT equilibration (T=300 K, P=1 bar) without position restraint and with the same coordination restraint as (iv).

(vi)	100 ns of NPT equilibration without position restraint and with reduced coordination restraint (1×10^4^ kJ mol^-1^ nm^-2^) for fully relaxing the whole system with stable Mg^2+^ coordination.

The last frame of the 100 ns NPT equilibration (vi) was used as the initial structure of all the four model systems. For each system, the model structure was subjected to the following relaxation with the same Zn^2+^ and Mg^2+^ coordination restraints as (vi):

(vii)	10,000 steps of energy minimization using steepest decent with position restraint (1×10^3^ kJ mol^-1^ nm^-2^) on the heavy atoms of RNA and the two catalytic Mg^2+^ ions;

(viii)	10,000 steps of energy minimization using steepest decent without position restraint;

(ix)	200 ps of NVT equilibration (T=300 K) with position restraint (1×10^3^ kJ mol^-1^ nm^-2^) on all the heavy atoms of protein/RNA, the structural Zn^2+^ ions and the two catalytic Mg^2+^ ions;

(x)	500 ps of NPT equilibration (T=300 K, P=1 bar) with the same restraints as (ix)

The final frame of NPT equilibration (x) was used as the starting structure of 10 replicas of 100 ns NPT production simulations (T=300 K, P=1 bar) with different initial velocities. Simulated annealing was performed in the first 2 ns with gradual temperature increase from 50 K to 300 K. The final 10 ns trajectories of each 100 ns replicas were used for analysis.

All the MD simulations were performed using GROMACS 2024.5.^[19]^ Integration time step of 2 fs was used. The system temperature was adjusted to 300 K by the V-rescale^[20]^ algorithm and pressure to 1 bar C-rescale^[21]^ algorithm, with relaxation time constant of 0.1 ps and 0.5 ps respectively. Thes system was partitioned into two temperature coupling groups: (1) Protein, RNA, two catalytic Mg^2+^ ions and five structural Zn^2+^ ions; (2) Water molecules, neutralizing Mg^2+^ and Cl^-^ ions. The initial velocities were assigned from the Maxwell-Boltzmann distribution at 300 K for equilibration runs and 50 K for production runs. For non-bonded cutoffs, neighbour list was set to 1.2 nm, and PME (rcoulomb=1.2 nm) and L-J potential (rvdw=1.2 nm and vdw-modifier=potential-shift) were used. All the h-bonds were constrained with SHAKE algorithm except during energy minimization.
